# Multivalent Nanobodies for Potent and Broad Neutralization of *Staphylococcus aureus* Toxins

**DOI:** 10.64898/2025.12.24.696403

**Authors:** Yong Joon (Jeffrey) Kim, Nicholas R. Walton, Wei Huang, Madison Lee, Yufei Xiang, Zhe Sang, Chaya Sussman, Sarah K.L. Moore, Derek J. Taylor, Kong Chen, Jaime L. Hook, John K. McCormick, Yi Shi

**Affiliations:** Center for Protein Engineering and Therapeutics, Department of Pharmacological Sciences, Icahn School of Medicine at Mount Sinai, New York, NY 10029, USA; Department of Microbiology and Immunology, University of Western Ontario, London, Ontario, N6A5C1, Canada; Department of Pharmacology, Case Western Reserve University, Cleveland, OH 44106, USA; Division of Pulmonary, Critical Care and Sleep Medicine, Department of Medicine, Icahn School of Medicine at Mount Sinai, New York, NY 10029, USA; Department of Microbiology, Icahn School of Medicine at Mount Sinai, New York, NY 10029, USA; Global Health and Emerging Pathogens Institute, Icahn School of Medicine at Mount Sinai, New York, NY 10029, USA; Stony Brook University, Stony Brook, NY 11794; Division of Pulmonary, Allergy, and Critical Care Medicine, Department of Medicine, University of Pittsburgh, Pittsburgh, PA 15213, USA

**Author notes:** Correspondence: Yi Shi.

## Abstract

*Staphylococcus aureus* is a leading cause of lethal bacteremia and pneumonia, which are driven by potent virulence factors such as T-cell superantigens and alpha hemolysin. *S. aureus* has among the highest rates of antibiotic resistance, yet no vaccines or alternative therapies are available despite decades of research. Here, we developed a repertoire of potent, high affinity nanobodies (Nbs) targeting key toxins in *S. aureus* infection, including superantigens (SAgs) SEB, SEC, TSST-1, and Hla. Comprehensive cryo-EM and AlphaFold3 analyses of these Nbs, which were elicited with clinical cocktail vaccines, revealed diverse neutralizing epitopes and mechanisms that provide strategic insights for immunotherapy and vaccine design. Guided by these findings, we engineered highly stable, multivalent, and multifunctional Nb constructs. These constructs included an aerosolizable trimeric Nb with enhanced neuralization activity against Hla and SEC, and an ultrapotent decameric Nb-IgG-Fc fusion construct against a wide range of major toxins in *S. aureus* sepsis (SEB, SEC, TSST-1, and Hla). These multifunctional Nbs demonstrated promising protective activity in murine models of pneumonia and sepsis, underscoring their potential as versatile immunotherapies that address the complex virulence profiles of *S. aureus*. Our work lays a foundation for precision immunotherapies beyond current treatment options to combat complex bacterial infections with multiple virulence mechanisms.

**Significance statement:** *S. aureus* is among the most common, antibiotic-resistant, and deadly causes of bacterial infections. We developed nanobodies against clinically significant virulence factors in *S. aureus* sepsis and pneumonia, including superantigens (SAgs) SEB, SEC, and TSST-1 as well as pore forming toxin Hla. These nanobodies displayed complete and potent neutralization of each toxin, exploiting a wide variety neutralizing mechanisms. Structural investigation of these diverse neutralizing nanobodies, which were elicited in llamas using clinically investigated cocktail vaccines, highlighted the importance of disrupting SAg interaction with TCR or MHCII and potential flaws in targeting poorly neutralizing conserved SAg epitopes using vaccine cocktails. Nb leads against each toxin were combined in different multivalent configurations, including an aerosolizable trimeric Nb and a half-life extended decameric Nb IgG Fc fusion construct. This work highlights multivalent nanobodies as a comprehensive yet therapeutically precise drug platform that addresses the complex virulence profiles of bacterial infectious diseases.

## Introduction

*Staphylococcus aureus* (*S. aureus*) is responsible for one-eighth of global bacterial infections (*1–3*) and colonizes 30–50% of healthy individuals (*4, 5*). In patients with weakened immunity, invasive procedures, or breached tissue barriers, *S. aureus* can cause severe infections including hospital-associated pneumonia, sepsis, osteomyelitis, and soft tissue infections (*6–9*). Recent global estimates attribute 1.1 million deaths annually to *S. aureus*, primarily due to pneumonia and bloodstream infections (*2*). The widespread emergence of methicillin-resistant *S. aureus* (MRSA) intensifies the challenge of treating these infections. MRSA is therefore designated an “Urgent Threat” by the CDC (*10*) and a “High Priority” pathogen by the WHO (*11–13*).

Despite their promise as alternatives to traditional antibiotics, vaccines and monoclonal antibodies (mAbs) against *S. aureus* have repeatedly failed clinical trials. *S. aureus* vaccines elicit titers of limited protective duration and efficacy (*14–16*). Monoclonal antibodies have also yet to display clinical benefit, likely because they target just one of many *S. aureus* virulence factors. These challenges underscore the need for alternative strategies that broadly and efficaciously neutralize key *S. aureus* pathogenic mechanisms.

T-cell superantigens (SAgs) and alpha hemolysin (Hla) are amongst the most critical *S. aureus* virulence factors that drive severe infections such as pneumonia and sepsis (*17, 18*). SAgs such as staphylococcal enterotoxin B (SEB), staphylococcal enterotoxin C (SEC), and toxic shock syndrome toxin-1 (TSST-1) are expressed in many clinical isolates (*19*) and these three particular SAgs have been historically associated with Staphylococcal toxic shock syndrome (*20*). Although *S. aureus* strains can encode many SAgs simultaneously in different combinations, TSST-1 as well as SEB and SEC have been strongly implicated in severe *S. aureus* infections (*21*). SEB, SEC, and TSST-1 are among the best described superantigens with strong implications in *S. aureus* diseases such as pneumonia and sepsis (*18, 22–31*). SAgs cross-link TCR Vβ domains with MHCII molecules on antigen-presenting cells to potently stimulate up to 30% of T cells, compared to ∼0.001-0.01% in normal adaptive responses. This SAg-driven hyperimmune response can induce cytokine storm mediated shock and drive infective endocarditis by compromising endothelial integrity and healing (*21, 32, 33*). SAgs also promote immune evasion through T cell exhaustion and conversion of follicular helper T cells into cytotoxic phenotypes that kill antibody producing MHCII^+^ B cells (*34*) (*35*).

Hla is a highly conserved pore-forming toxin with cytotoxic and inflammatory activities (*32, 36–38*), and its central role in pneumonia and sepsis has been thoroughly established (*36, 39–44*) (*45*). Secreted as a monomer, Hla oligomerizes into cytotoxic heptameric pores on host cell membranes (*46*). Hla drives complicated infections and overall morbidity by damaging immune and epithelial cells, promoting systemic bacterial dissemination, and inducing thrombotic multi-organ damage through systemic platelet aggregation (*40, 47*). Hla synergizes with SAgs with distinct inflammatory pathways to drive inflammatory and cytotoxic tissue damage and vascular leakage, promoting sepsis and systemic infection (*48, 49*).

Given the significant clinical burden of *S. aureus* and the limitations of current therapeutics, alternative treatments are urgently needed. Nanobodies (Nbs), which are antigen-binding domains from camelid heavy chain-only antibodies, represent a new class of antibody therapeutics. Nbs are minimal, monomeric antibody fragments (∼15kDa) that display excellent target specificity, solubility, and stability (*50*). Their monomeric nature and marked stability make Nbs highly amenable for engineering, including the development of multispecific constructs with expanded functional activities (*51, 52*). Nbs can target diverse epitopes including small and cryptic sites that are not readily accessible with mAbs (*53–58*). Nb-based therapies have been clinically approved, displaying excellent safety profiles (*59–62*).

Previous efforts have identified Nbs against specific S. aureus antigens; however, most exhibit limited neutralization efficacy, reflecting the marked bioactivity of these toxins and the challenges inherent in identifying effective, potent neutralizers(*63–65*)(*66*)(*67*).Here, we report the development of high-affinity and potently neutralizing Nbs against a panel of key *S. aureus* virulence factors including both SAgs (SEB, SEC, TSST-1) and Hla. Structural characterization of Nb-toxin complexes revealed diverse epitopes and neutralization mechanisms, offering insights into immunotherapy and vaccine design. To address the multifaceted nature of *S. aureus* infections, we further engineered multivalent, stable, aerosolizable, and half-life extended Nb constructs simultaneously targeting these four major virulence factors. These ultrapotent constructs demonstrated protective efficacy in two murine models, underscoring their potential as a novel therapeutic platform against complex *S. aureus* infections.

## Results

### Discovery and functional characterization of SAg Nb repertories

To raise high-quality Nbs, llamas were immunized (**Figure 1A**) using clinically investigated vaccine toxoids of SEB, SEC, TSST-1 and Hla (**Methods**), each containing mutations to abrogate Hla oligomerization (*68–70*) or SAg interaction with MHCII and TCR (*71–76*). Specific antibody titers against wild-type SEB, SEC, TSST-1, and Hla were observed after immunization, with ∼2-log titer enhancements compared to pre-immunization sera (**Figure 1B**). Antigen-specific, full-length heavy-chain-only antibodies (hcAbs) were isolated and digested with trypsin/LysC or chymotrypsin for mass spectrometry-based identification(**Figure 1A**) (*53*). LC-MS spectra of the digested full-length hcAb peptides were searched against a PBMC-derived variable domains of a heavy chain-only antibody (VHH) genomic database. Candidate peptide-spectrum matches were subsequently analyzed using our published software, *AugurLlama* (*53*), which effectively filters false positives and enables confident assignment of CDR peptides, particularly CDR3, allowing deconvolution and identification of hundreds of diverse Nb CDR3 sequences against wild-type SEB, SEC, TSST-1, and Hla (**Figure 1C-E**).

**Figure 1.**
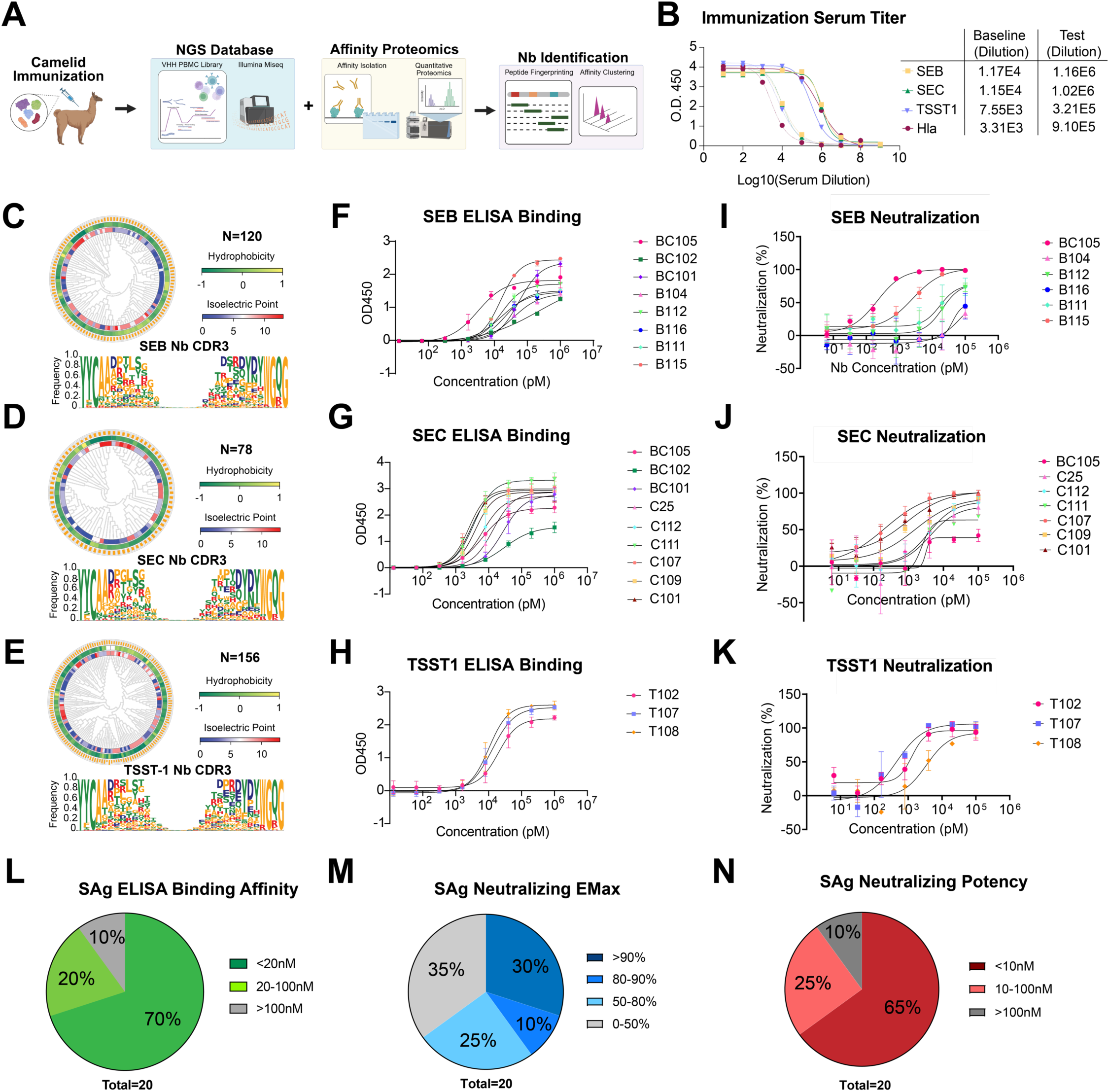
Discovery of Nbs against SEB, SEC, and TSST1. (**A**) Schematic of Nb discovery pipeline. (**B**) Serum titers before and after immunization. (**C-E**) Identified Nb CDR3 diversity against SEB (C), SEC (D), and TSST-1 (E). (**F-H**) ELISA binding for Nbs against SEB (F), SEC (G), and TSST-1 (H). ELISA experiments completed in triplicate and fitted according to 4PL regression. (**I-K**) Human PBMC assays for functional neutralization of SEB (I), SEC (J), and TSST-1 (K). PBMC experiments completed in triplicate and fitted according to 4PL regression. (**L-N**) Summary of all anti-SAg Nbs for (L) ELISA binding EC50s (M) Maximum neutralizing efficacy and (N) Neutralizing IC50s.

A sequence-diverse panel of Nbs against SEB, SEC, and TSST-1 were expressed and purified from *E. coli*. ELISA confirmed high-affinity binding to all three SAgs (**Figure 1F–H** & **Table 1**). Eight Nbs were identified for SEB, nine for SEC, and three for TSST-1. Among SEB Nbs, BC105 and B115 exhibited the highest affinities (EC50s of 3.2 nM and 3.7 nM, respectively). Six of nine SEC Nbs displayed single-digit nM EC50s, while TSST-1 Nbs showed EC50s in the 10–20 nM range. Some Nbs (BC101, BC102, and BC105) displayed binding against both SEB and SEC, which have high sequence homology (∼65%) (**Supplemental Figure 1A**). BC105 bound strongly to both SEB (EC50 = 3.2 nM) and SEC (EC50 = 6.3 nM), whereas BC101 and BC102 showed moderate binding activities.

**Table 1.**
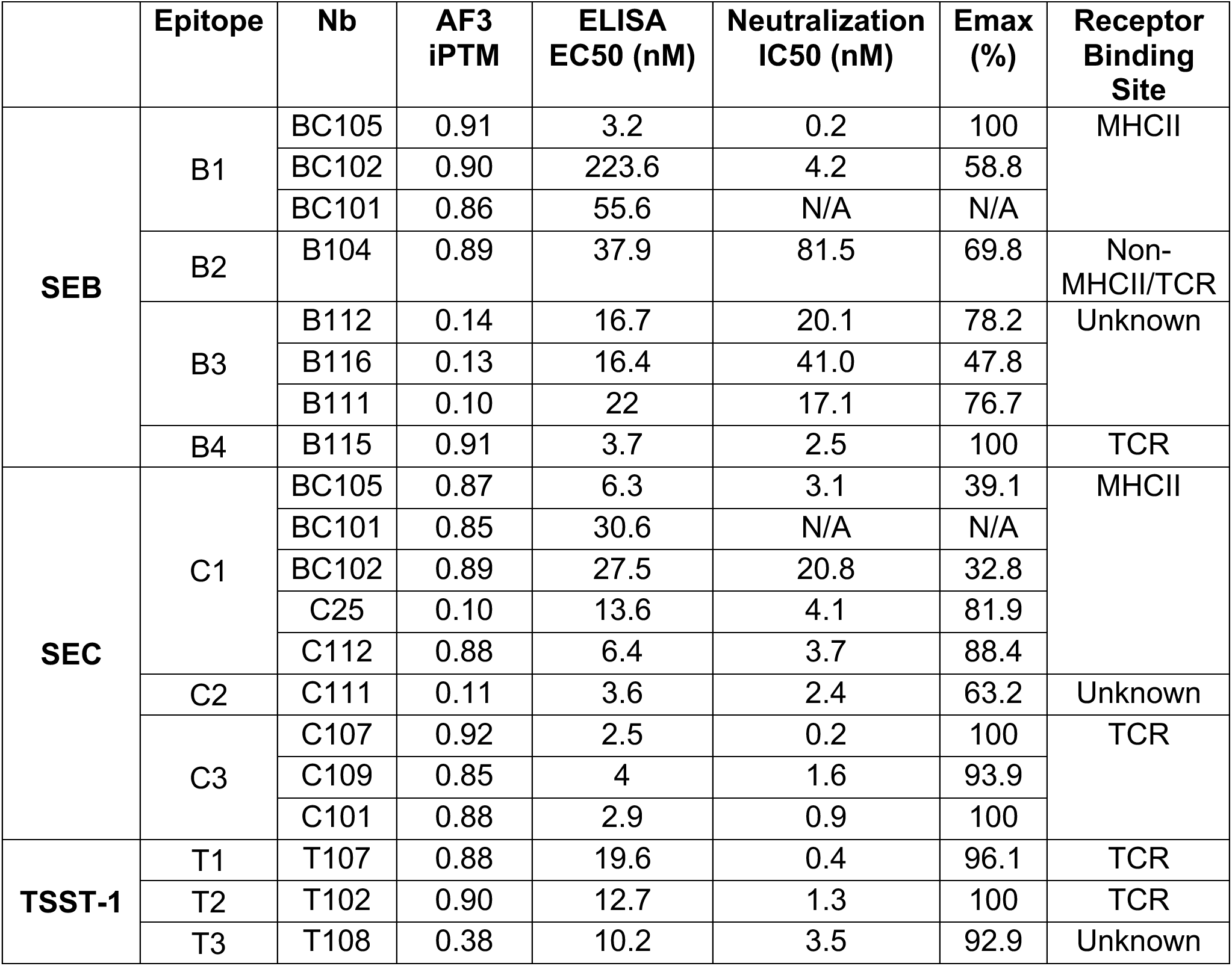
Summary of epitope classification, ELISA binding, neutralization IC50, and neutralization efficacy for all Nb hits against superantigens SEB, SEC, and TSST-1.

Neutralization activities of anti-SAg Nbs were assessed *in vitro* using human PBMC stimulation assays (**Methods**). Lead Nb B115 completely neutralized SEB with high potency (IC50 = 2.5 nM) (**Figure 1I**). The majority of anti-SEC Nbs (5/9) displayed strong neutralizing efficacy and nM or better potencies, with lead Nb C107 neutralizing SEC with a potency of 211 pM (**Figure 1J**). All three anti-TSST-1 Nbs completely neutralized TSST-1 with single digit nM or better neutralizing potencies, with lead T107 neutralizing at a potency of 445 pM (**Figure 1K**). Cross-reactive Nbs (BC105, BC101, BC102) had notably inconsistent or poor neutralizing profiles. BC105 fully neutralized SEB (IC50 = 0.2 nM) but showed lower efficacy and potency against SEC (Emax = 39.1%, IC50 = 3.1 nM). SEC-specific Nbs displayed significantly higher neutralizing efficacies than cross-reactive Nbs, and a similar trend was observed for SEB-specific Nbs (**Supplemental Figure 1B-D**). Cross-reactive Nbs may target conserved residues that do not overlap with key SAg functional sites required for neutralization.

Next, Nbs were classified by epitope through competitive size exclusion chromatography analyses of different Nb-toxin combinations (**Methods**) (*54*). Our analysis revealed multiple non-overlapping epitopes against each SAg, identifying 4 epitopes for SEB, 3 epitopes for SEC, and 3 epitopes for TSST-1 (**Supplemental Figure 2** & **Table 1**). SAg neutralization was found to be highly toxin and epitope dependent, with only select epitopes showing consistently efficacious neutralization activity (**Supplemental Figure 3**). Notably, cross-reactive Nbs such as BC105, which displayed variable neutralization activities, all targeted a single conserved epitope shared by SEB and SEC. In contrast, Nbs against certain epitopes specific to individual SAgs such as B115 (SEB epitope B4), C107 (SEC epitope C3), and T107 (TSST-1 epitope T1) showed strong neutralization. This data highlights the importance of selecting Nbs that engage key functional SAg epitopes, which do not show strong SAg cross-reactivity, for robust neutralizing activities.

### Structural insights in Nb neutralization

To understand the neutralization mechanisms of anti-SAg Nbs, we determined a 3.1 Å cryo-electron microscopy (cryo-EM) structure of a trimeric complex of SEC with two potent Nbs, C107 and C112 (**Figure 2A**). Nb C107, from the highly neutralizing C3 epitope class, binds directly to the TCR binding interface on SEC using its CDR3, FR2, and FR3 regions. C107 buries deeply within a cleft on SEC formed by its N-terminal beta barrel and helix 2 (**Figure 2B-D**). Key stabilizing interactions include GLU104 (CDR3) with residues ASN50 and THR47 of SEC, and ARG53 (CDR2) with SEC ASP57 (**Figure 2E**).

**Figure 2.**
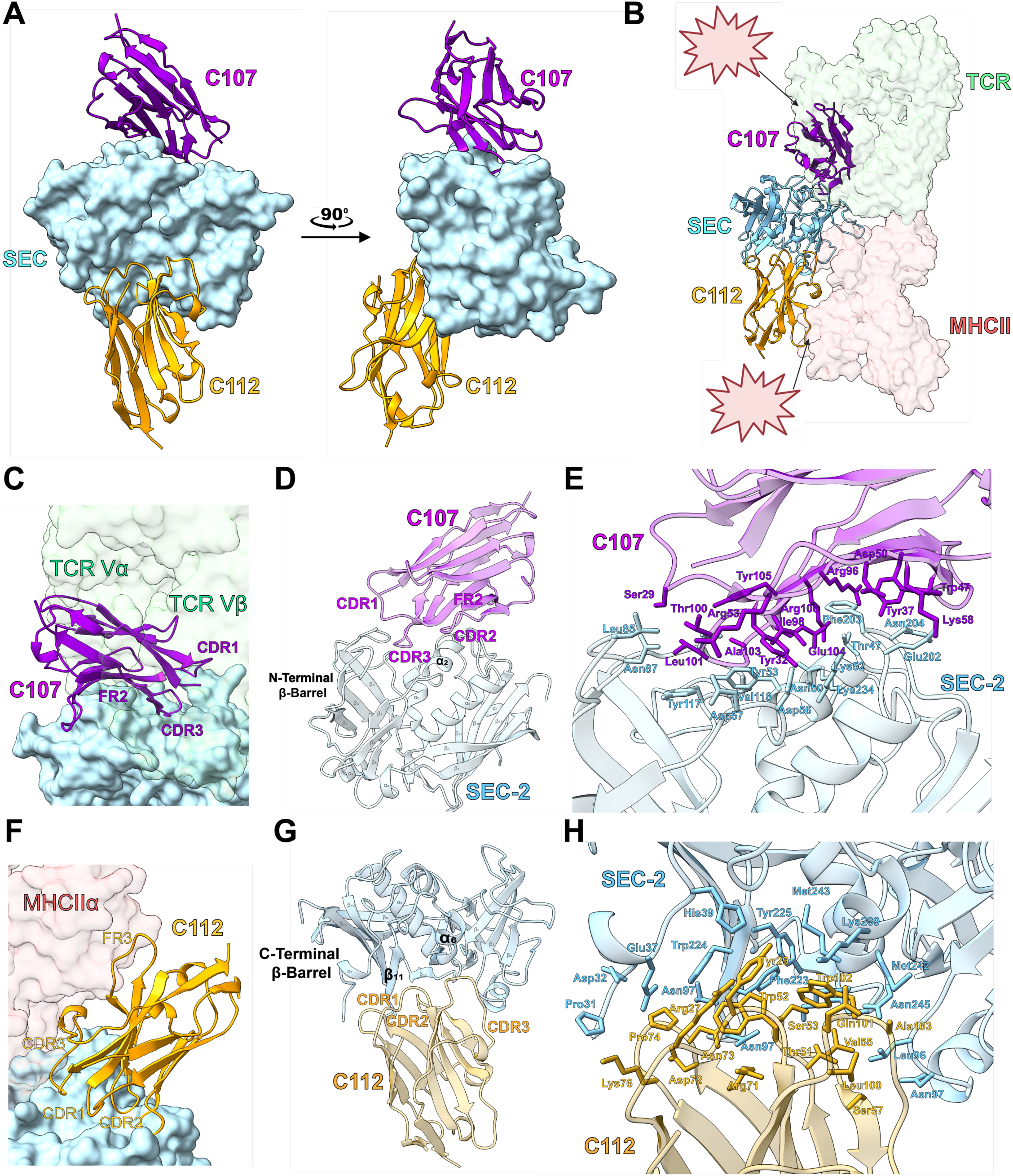
**(A**) Cryo-EM structure of Nb C107 and C112 in complex with SEC-2 (Overall resolution=3.1 angstroms). (**B**) Nb-SEC structure superimposed with co-complex structures of SEC with TCR (PDB: 1JCK) and MHCII (1JWM). The fully assembled TCR from PDB:4C56 was used to fully visualize TCR densities (**C**) Zoomed view of Nb C107 and its overlap with TCR. (**D**) C107-SEC secondary structure interactions. (**E**) C107-SEC residue interactions. (**F**) Zoomed view of Nb C112 and its overlap with MHCII. (**G**) C112-SEC secondary structure interactions. (**H**) C112-SEC residue interactions.

Nb C112 targets a distinct concave epitope of SEC formed by the C-terminal beta barrel and helix 6, located adjacent to the MHCII binding site (**Figure 2F-G**). CDR3 and FR3 of C112 sterically clash with the MHCII DR alpha binding domain, explaining its potent neutralization activity (**Figure 2B&F**). Stabilizing interactions include TYR29 (CDR1) with SEC ASP226, as well as SER53 (CDR2) with SEC ASP246 (**Figure 2H**). Overall, cryo-EM structure analyses reveal that direct TCR competition, as seen with C107, was crucial for neutralization. In contrast, the activity of epitope C1 Nbs, which includes C112 as well as SEB and SEC cross-reactive Nbs, likely depends on variable binding angles and indirect steric overlaps with MHCII.

To expand these insights across other anti-SAg Nbs, we used AlphaFold 3 (AF3) to model Nbs targeting SEB, SEC, and TSST-1 (**Figure 3, Supplemental Figures 4–6**)(*77*). High-confidence AF3 models (**Methods**) closely matched cryo-EM and epitope binning data (**Supplemental Figure 4**). For SEC, seven of nine Nbs were confidently predicted. Epitope C3 Nbs (C101, C107, C109) consistently occupied the TCR binding site, correlating with strong neutralization (**Figure 3B**). Epitope C1 Nbs showed variable competition with MHCII depending on framework orientation (**Supplemental Figure 4**). Nb BC101, which is oriented away from MHCII, displayed notably weak neutralizing activity. Similar patterns were seen for SEB, where five of nine Nbs were confidently modeled, with lead Nb B115 engaging the TCR site directly and epitope B1 Nbs binding to an epitope adjacent to the MHCII binding site with variable degree of overlap with MHCII (**Supplemental Figure 5**). For TSST-1, AF3 confidently predicted two Nbs (T102, T107), both targeting distinct regions of the TCR binding site (**Figure 3C** & **Supplemental Figure 6**), consistent with their potent neutralization. Overall, Nbs were found to neutralize SAgs by disrupting SAg interaction with either TCR or MHCII, with inhibition of TCR binding showing the most consistent and complete neutralizing efficacy.

**Figure 3.**
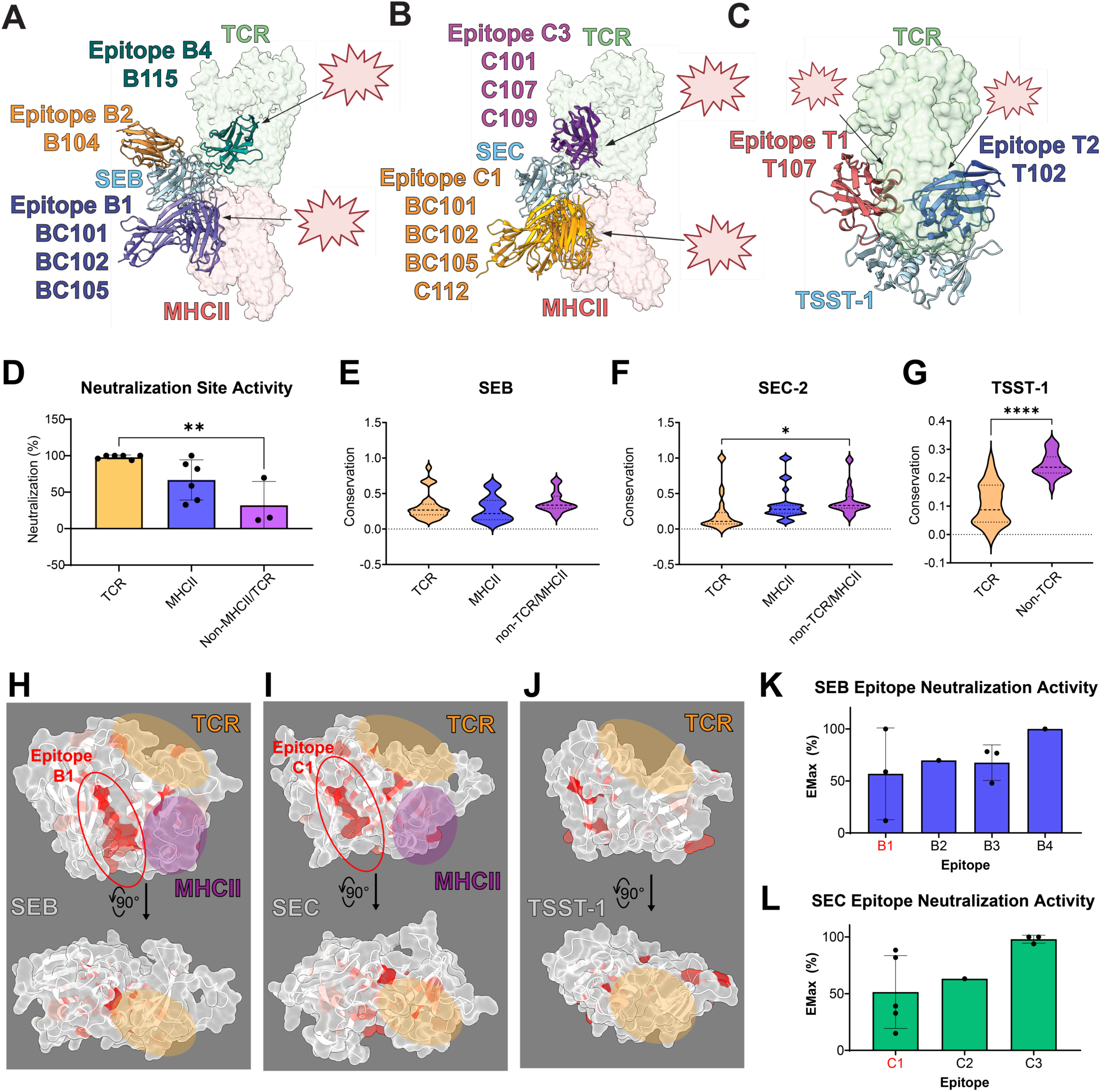
High confidence AF3 models for Nbs against **(A**) SEB, (**B**) SEC-2, and (**C**) TSST-1. (**D**) Neutralizing activities of anti-SAg Nbs categorized by TCR or MHCII competition. Comparisons made using ordinary one-way ANOVA analysis (* is p<0.05 and ** is p<0.01) (**E-G**) Degree of sequence conservation of TCR/MHCII receptor binding sites among all known *S. aureus* superantigens. Comparisons made using ordinary one-way ANOVA analysis (* is p<0.05 and ** is p<0.01) (**H-J**) Residue conservation score plotted as red on structures of SEB, SEC, and TSST-1. TCR and MHCII binding regions highlighted in yellow and purple, respectively. Conserved Nb epitopes B1 and C1 circled in red. (**K-L**) Comparison of Nb neutralizing EMax of conserved epitopes B1 and C1 compared to other epitopes.

These Nbs are representative of antibody repertoires elicited by current clinical vaccine formulations of SAg toxoid cocktails (*78–82*), which are used to elicit antibodies against epitopes conserved across the 26 different *S. aureus* SAgs. Our repertoire of high quality Nbs, encompassing 10 different SAg epitopes, was used to assess the localization and neutralizing efficacy of conserved epitopes elicited with cocktail vaccines. Mapping residue conservation across 26 *S. aureus* SAgs revealed that TCR and MHCII binding sites, which were necessary for effective toxin neutralization (**Figure 3D**), are less conserved than other surface regions (**Figure 3E–G**). This heterogeneity in the TCR and MHCII binding sites is thought to have evolved from *S. aureus* targeting different TCR and MHCII subtypes to target larger proportions of immune cell populations (*83*) (*18, 20, 84*). For example, SEC’s TCR binding site (0.22 ±0.27) was significantly less conserved than non-functional areas (0.40 ± 0.21). Cross-reactive Nbs (Epitopes B1 and C1) bound conserved interfaces proximal to the MHCII site shared across SAgs (**Figure 3H-I**), but still displayed poor overall SAg cross-reactivity (**Supplemental Figure 7**) and inconsistent neutralization activities (**Figure 3K-L**). Structure-function analyses of our Nbs, which include Nbs targeting the most conserved SAg residues, indicate that eliciting a protective broad-spectrum antibody response with current cocktail vaccines is improbable given the particularly low conservation of therapeutically relevant TCR/MHCII binding sites and poor overall sequence homology among different SAgs.

### Discovery and characterization of anti-Hla Nbs

Two high-affinity lead nanobodies (Nbs), H204 and H216, were identified against recombinant WT alpha hemolysin (Hla), with ELISA EC50 values of 1.1 nM and 9.3 nM, respectively (**Figure 4A**). Epitope binning by size exclusion chromatography showed that they bind distinct epitopes on Hla (**Supplemental Figure 2**). In rabbit blood hemolysis assays, both Nbs conferred complete protection against Hla activity at potencies of 1.7 nM for H204 and 2.8 nM for H216 (**Methods, Figure 4B**). High-quality AF3 models showed that H204 binds the beta-sandwich “cap” region of the Hla monomer (**Figure 4C-E**), where it sterically clashes with adjacent Hla subunits to potentially block pore oligomerization. H204 interacts through its CDR3 loop with Hla loops L183–K190 and P129–K136. In contrast, H216 binds the inactive conformation of the “stem” region (residues E137–Y174) (**Figure 4F**), which otherwise forms the transmembrane beta barrel pore to mediate osmotic cytotoxic effects (*36*).

**Figure 4.**
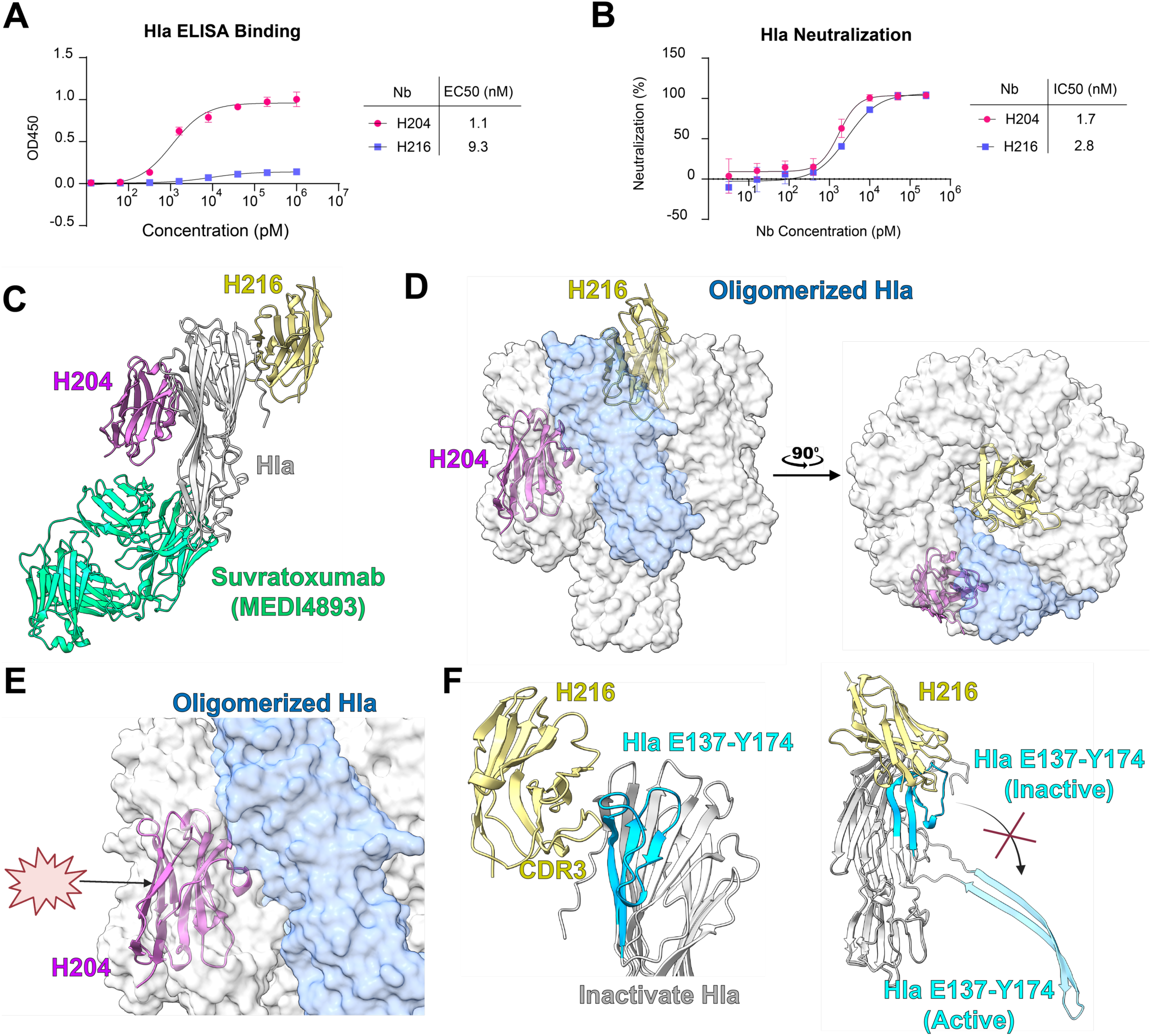
**(A**) ELISA binding curves of H204 and H216 against Hla WT (n=3). Neutralization curves fitted using 4PL regression analyses. (**B**) Neutralizing activities of H204 and H216 against recombinant Hla in rabbit red blood cell hemolysis assays (n=3). Neutralization curves fitted using 4PL regression analyses. **(C**) AF3 models of H204 (iPTM: 0.83) and H216 (iPTM: 0.86) complexed with Hla, superimposed with structures of Suvratoxumab-Hla co-complex (PDB: 4U6V). (**D**) AF3 models of H204 and H216 superimposed with assembled Hla structure (PDB: 7AHL). (**E**) Zoomed in view of H204 binding in assembled Hla state. H204 sterically clashes with adjacent Hla subunit. (**F**) Zoomed in view of H216 making contacts with the Hla stem region E137-Y174.

### Multivalent Nb engineering against SEC and Hla

Guided by structural insights, we engineered two multispecific Nbs: a biparatopic dimer (C112-C107) and a trivalent construct (C112-C107-H204). These constructs were designed to enhance neutralization potency against SEC, and C112-C107-H204 had additional toxin specificity against Hla (**Figure 5A-B**). SEC was chosen due to its critical role in bacteremia, pneumonia, and infective endocarditis, (*21, 29, 85, 86*) as well as the availability of highly neutralizing Nbs against different SEC epitopes. Hla was incorporated into the trivalent design to provide broader protection, given its high conservation across *S. aureus* strains and synergistic pathogenicity with SAgs in necrotizing pneumonia and sepsis. A flexible GGGGSx5 peptide linker was inserted between Nbs to enhance avidity effects.

**Figure 5.**
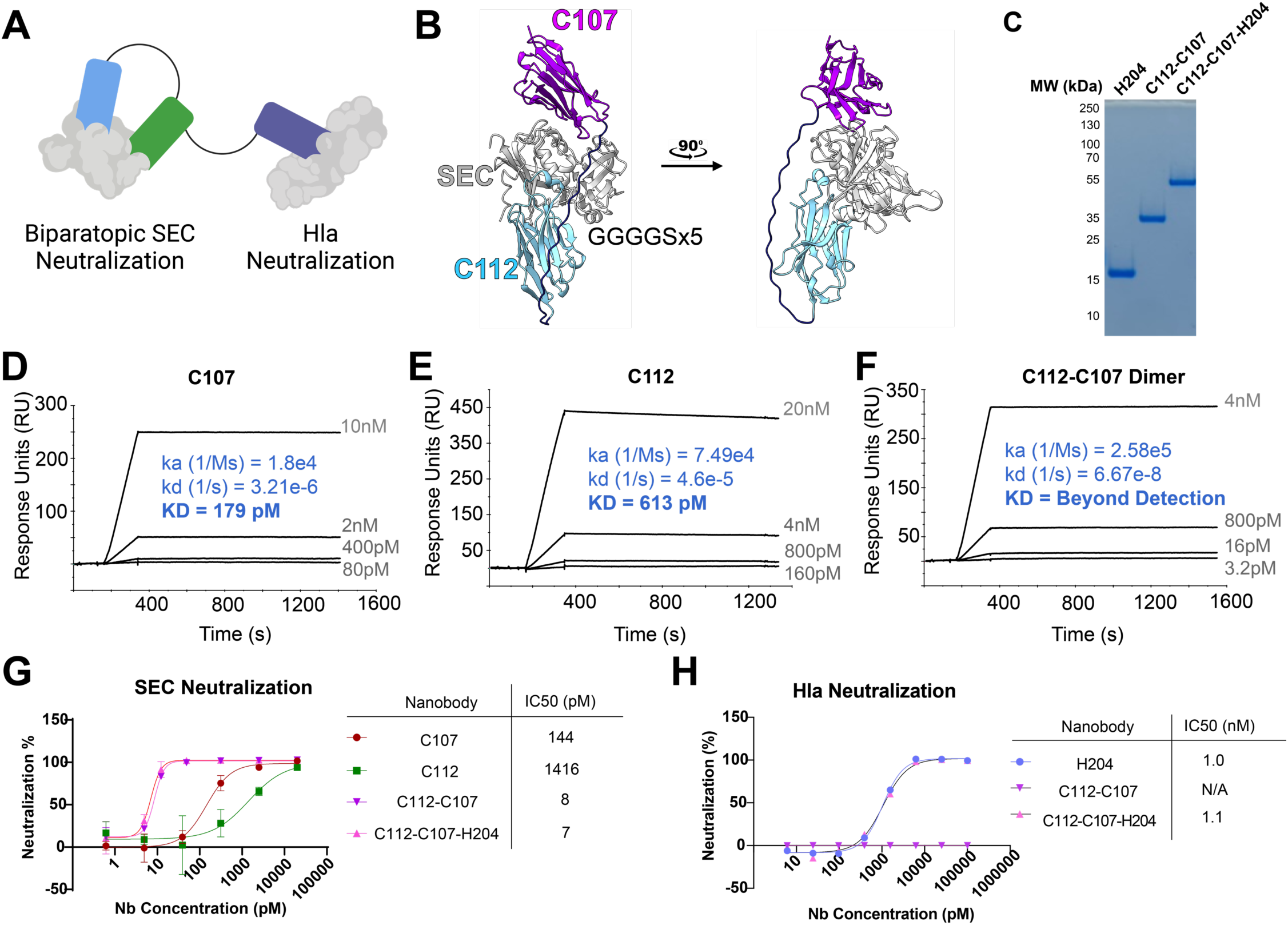
Trivalent Nb against SEC and Hla. **(A**) Cartoon schematic of trimeric Nb with biparatopic binding of SEC and the addition of an anti-Hla lead (**B**) High confidence AF3 model of C112-C107 bound to SEC (**C**) SDS-PAGE of H204 monomer, C112-C107 dimer, and C112-C107-H204 trimer (**D-F**) SPR measurements of Nb C107, C112, and C112-C107. Each analyte concentration measured in duplicate. (**G**) Neutralization of SEC in PBMC assays by multivalent Nbs compared to monomeric leads (n=3) (**H**) Neutralization of Hla by C112-C107-H204 compared to H204 (n=3). All neutralization curves analyzed using 4PL regression analyses.

Both constructs were successfully cloned, expressed, and purified with high yields (**Figure 5C**). Surface Plasmon Resonance (SPR) analysis showed that while monomeric Nbs C112 and C107 displayed K_D_’s of 179 pM and 600 pM, respectively (**Figure 5D-E**), the biparatopic C112-C107 exhibited strong avidity effect to SEC, with binding affinity beyond the detection limit of SPR (**Figure 5F**). The ultrahigh-affinity and enhanced avidity of these Nbs were independently confirmed using BioLayer Interferometry (BLI) (**Supplemental Figure 8**).

Dissociation of these Nbs was extremely slow, precluding accurate determination of koff values; therefore, the reported affinities should be interpreted as conservative lower-bound estimates. Nevertheless, the kinetic behaviors observed by SPR and BLI were consistent with one another and correlated with the low-picomolar functional potencies measured *in vitro*. The constructs also demonstrated high structural and functional stability after aerosolization (**Supplemental Figure 9**), supporting their feasibility for aerosolized delivery in pulmonary infections (*87, 88*).

C112-C107 exhibited an 18-fold increase in SEC neutralization potency in PBMC stimulation assays, achieving a low picomolar IC50 of 8 pM (**Figure 5G**). Incorporation of H204 in the trivalent construct maintained enhanced SEC neutralization potency (IC50 = 7 pM) and conferred potent anti-Hla activity (**Figure 5H**). Our results demonstrate high engineering potential and developability of affinity-matured Nbs (*51*), offering a promising therapeutic strategy against multifactorial *S. aureus* infections.

### Protective efficacy of a trivalent Nb in murine models of sepsis and pneumonia

To assess *in vivo* efficacy, we tested trivalent Nb C112-C107-H204 in two infection models. In a murine sepsis model (**Figure 6A**), we used humanized HLA-DR4-IE (DRB1*0401) mice (n=10) infected intravenously with the MRSA strain MW2, which expresses SEC and Hla as key virulence factors. The MW2 strain expresses SAgs (SEA, SEC, SEH, SEK, SEL, SEQ), Hla, and other exotoxins (PVL, LukED, LukGH, HlgAB, HlgCB). Transgenic HLA-DR4 mouse more faithfully models human responses to different *S. aureus* superantigens, which are adapted to human MHCII and TCR receptors (*84, 89, 90*). Transgenic mice with humanized MHCII receptors are proven models for delineating mechanisms of SAg-driven *S. aureus* disease and therapeutic effects of SAg downregulation or neutralization (*84, 91–96*).

**Figure 6.**
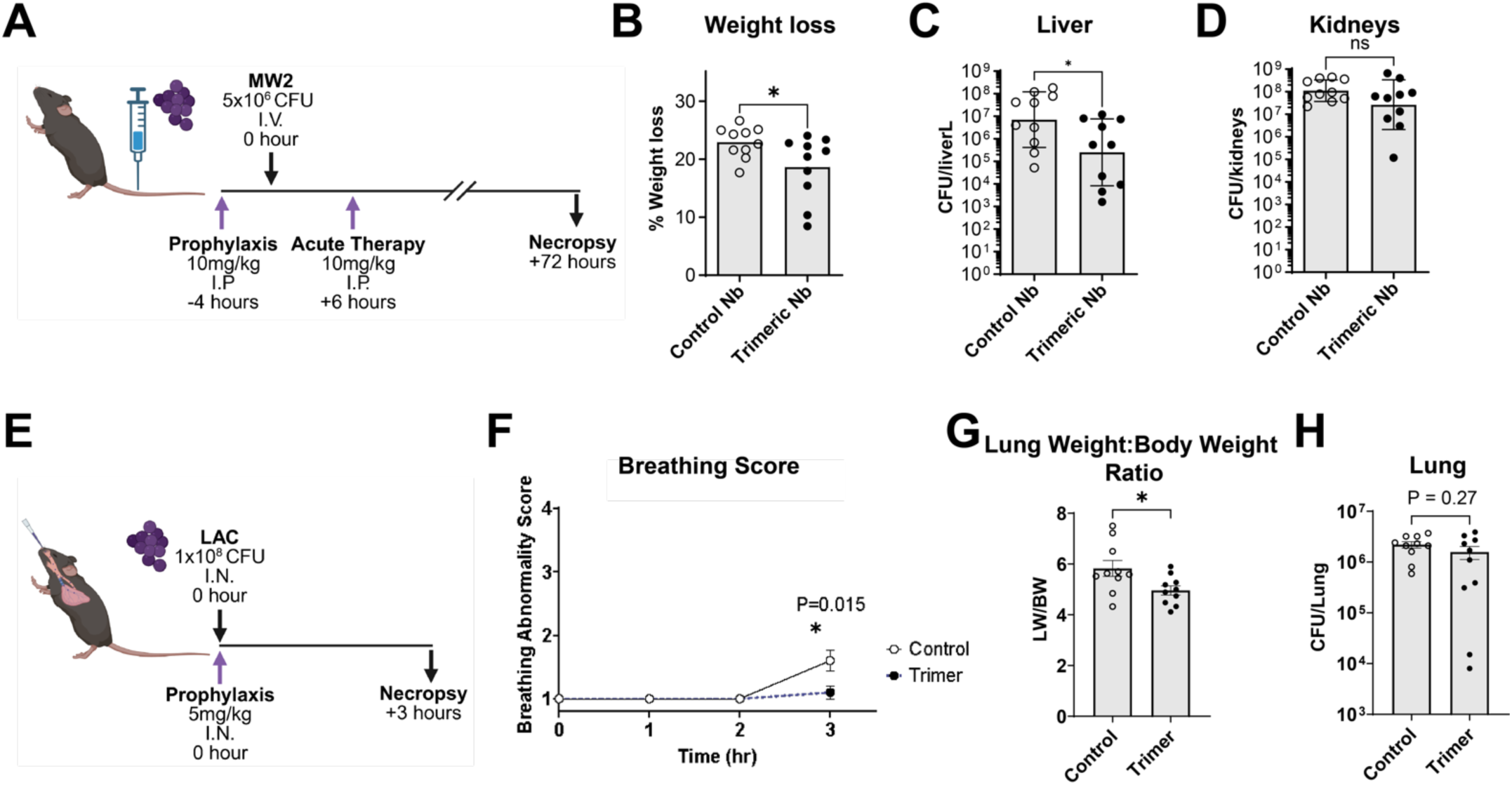
(**A-D**)Transgenic HLA DR4-B6 bacteremia models for assessing prophylactic treatment with C112-C107-H204 against a control Nb. (**A**) Experimental design *in vivo* bacteremia models. (**B**) Weight loss protection by Nbs. Statistical comparison calculated using unpaired t test (* is p<0.05) (**C**) Liver CFU protection by Nbs. Statistical comparison calculated using Mann-Whitney test (* is p<0.05) (**D**) Kidney CFU count after Nb treatment. Statistical comparison calculated using Mann-Whitney test (* is p<0.05) (**E-H**) WT Swiss-Webster mouse models of pneumonia. (**E**) Experimental design and timeline for pneumonia *in vivo* models. (**F**) Breathing score progression. Statistical comparison calculated using unpaired t tests (* is p<0.05) (**G**) Lung weight to body weight ratio (LW/BW) as measure of pulmonary edema. Statistical comparison calculated using unpaired t tests (* is p<0.05) (**H**) Lung CFU count after Nb treatment. Statistical comparison calculated using unpaired t tests (* is p<0.05).

Previous work showed that SEC deletion reduces sepsis-associated liver CFU by 1-2 logs in SEC expressing strain MW2 (*84*). Compared with the scrambled Nb control, prophylactic treatment with C112-C107-H204 significantly protected against MW2-induced weight loss (**Figure 6B**), indicating protection against overall bacteremia morbidity. Nb treatment also reduced liver bacterial burden by 15-fold compared to negative control Nb treatment (**Figure 6C**), indicating enhanced immune clearance and disruption of SEC-driven infection pathogenesis. Kidney CFUs were not significantly reduced (**Figure 6D**), consistent with outcomes in SEC deletion studies (*84*).

To evaluate the therapeutic potential of Hla neutralization in pneumonia, we used our established mouse model of acute staphylococcal lung infection (*97*). Although MW2 expresses Hla and transgenic HLA-DR4 mice are likely to maintain susceptibility to Hla, we sought to confirm the Hla neutralizing effects of C112-C107-H204 in a dedicated, validated model of Hla-driven infection. We intranasally instilled WT Swiss-Webster mice with LAC (1×10⁸ CFU), a USA300 MRSA strain that is prototypically associated with Hla driven lung infection (*98–100*), and the trivalent Nb to simulate nebulized Nb treatment (**Figure 6E**).

Experimental outcomes included breathing score, lung bacterial burden, and lung weight to body weight ratios as measures of pulmonary edema (*45, 97*). Breathing score, which is a composite measure of respiratory rate, rhythm, and effort (*97*), was monitored for 3 hours. The breathing score has been well established to correlate with bronchoalveolar lavage content of total protein, a direct measure of pulmonary edema and defining characteristic of acute lung injury (*97*). Lung edema is a hallmark direct marker of lung damage and inflammation, and lung weight to body weight ratio is a gold standard readout of lung edema (*97*).

Compared to the scrambled Nb control, C112-C107-H204 significantly improved breathing scores (n=10) (**Figure 6F**) and reduced lung weight-to-body weight ratios (**Figure 6G**), indicating that C112-C107-H204 decreased lung edema and inflammation. No significant lung CFU reduction, a readout expected as a delayed effect of Hla neutralization from enhanced immune clearance (*38*) (*101*), was observed given the short experimental duration of 3 hours (**Figure 6H**).

To further evaluate feasibility of aerosolized treatment using Nb C112-C107-H204, the construct was aerosolized using a mesh nebulizer and assessed for structural and functional integrity (**Supplemental Figure 9**). Nb C112-C107-H204 was able to withstand aerosolization without aggregation or major loss (**Supplemental Figure 9A**). Nb C112-C107-H204 also maintained robust binding activities against SEC and Hla after aerosolization (**Supplemental Figure 9B-C**).

Collectively, our data suggests that the trivalent Nb C112-C107-H204 simultaneously neutralizes SEC and Hla *in vivo* against 2 different strains.

### Engineering of Avidity- and Half-life-Enhanced Multifunctional Nb-Fc Construct Against Four Major Toxins

Building on the success of Nb trimer (C112-C107-H204) and as a proof-of-concept of expanded specificity and functionality, we engineered a decameric Nb-Fc fusion construct targeting SEB, SEC, TSST-1, and Hla (**Figure 7A**).

**Figure 7.**
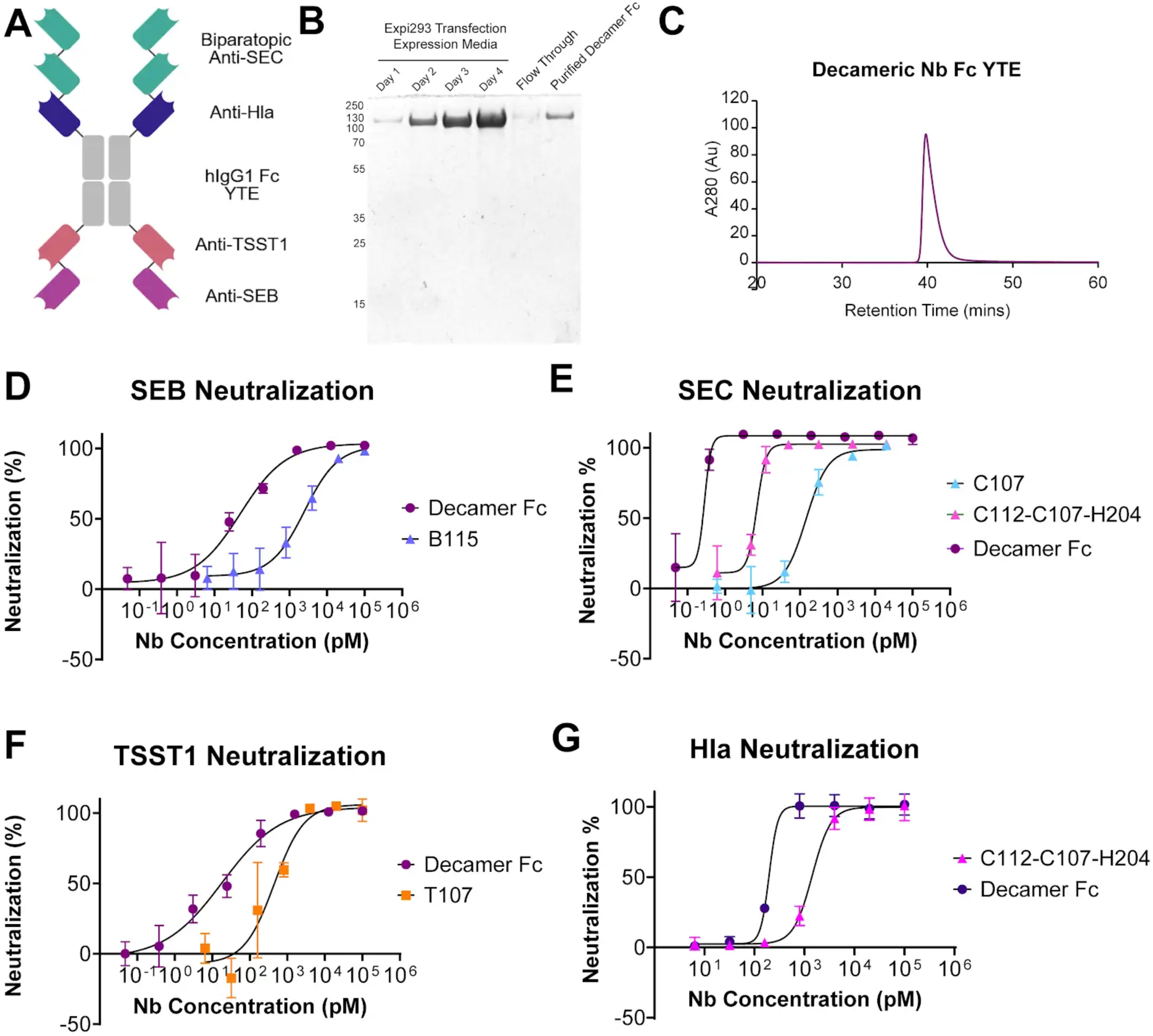
**(A**) Cartoon schematic of decameric Nb Fc YTE fusion construct against SEB, SEC, TSST-1, and Hla. Molecular weight of fully assembled construct is 200.5 kDa. (**B**) SDS-PAGE and (**C**) size exclusion chromatography profile of decameric Nb Fc fusion construct. *In vitro* neutralization activity of decameric Nb Fc YTE construct compared to respective lead Nbs against (**D**) SEB, (**E**) SEC, (**F**) TSST-1, (**G**) Hla. All neutralization curves generated in triplicate and fitted according to 4PL regression analyses.

We used an Fc YTE variant (M252Y, S254T, T256E) to enhance *in vivo* half-life (∼85 days in clinical trials (*102*)) while maximizing avidity enhancing effects. This design incorporated biparatopic anti-SEC Nbs (C112, C107) and anti-Hla Nb H204 into the N-terminus of the Fc, with additional lead Nbs against SEB (B115) and TSST-1 (T107) linked at the C-terminus using flexible GGGGSx5 linkers (**Figure 7A**). The addition of anti-SEB and anti-TSST-1 Nbs expands coverage against more superantigens, potentially providing wider coverage against different SAg producing *S. aureus* strains. The resulting decameric construct (MW = 200.5 kDa) was highly soluble and can be abundantly produced (∼ 0.5mg/ml of culture) in Expi293F cells (**Methods**, **Figure 7B, Supplemental Figure 10**) and showed strong, specific binding to all four toxins in ELISA binding assays (**Supplemental Figure 11**).

Functional *in vitro* assays demonstrated pM to sub-pM neutralization across all toxins. Notably, SEB neutralization improved 24-fold compared to B115 monomers (IC50=103 pM vs. 2.5 nM) (**Figure 7D**), while TSST-1 neutralization improved from 445 pM to 18 pM (**Figure 7F**). Hla neutralization also improved 9-fold (1.7 nM to 0.2 nM) (**Figure 7G**). SEC neutralization reached femtomolar potencies, yielding ∼21-fold enhancement over the trimer C112-C107-H204 and ∼480-fold enhancement over lead C107 (**Figure 7E**).

The decameric Nb Fc fusion construct also displayed promising neutralizing activities *in vitro* against supernatants of prototypical strains that express SEB (COL) and TSST-1 (MN8) (**Supplemental Figure 12**). The multivalent construct trended towards neutralization of COL supernatant in PBMCs comparable to SEB deletion (**Supplemental Figure 12A**) and displayed significant neutralization of MN8 supernatant in PBMCs comparable to TSST-1 deletion (**Supplemental Figure 12B**).

## Discussion

*S. aureus* employs multiple exotoxins to drive severe infections and undermine the development of protective immunity (*21, 48, 103*). The emergence of antimicrobial resistance, along with poor tissue penetration and adverse side effects of antibiotics, has limited the effectiveness of existing antibiotics (*2, 3, 11, 12*). Small molecule antibiotic development has stalled, and new drugs are typically derived from existing antibiotic classes. Vaccine efforts have also been unsuccessful, as pre-existing non-protective antibodies are reinforced by vaccination and interfere with the development of effective antibody responses (*15, 16*). While passive immunization with neutralizing mAbs can address some of these challenges, the multifactorial virulence of *S. aureus* demands therapies that neutralize multiple toxins. Nbs, with their small size, high epitope diversity, and engineering versatility, represent a promising therapeutic approach to overcome these limitations (*59, 60*).

Here, we successfully generated Nbs against clinically important *S. aureus* toxins in bacterial sepsis and pneumonia using a proteomics based Nb discovery pipeline (refs). This diverse panel of Nbs targeted multiple neutralizing epitopes and employed a wide range of neutralizing mechanisms of action. Structural investigations by cryo-EM and AF3 modeling revealed that Nbs that engage SAgs at their TCR binding sites exhibited superior neutralization activities compared to those targeting MHCII or non-TCR/MHCII regions. Structural analyses of these Nbs, which were elicited using vaccine cocktail approaches that have failed clinical trials, revealed that the most conserved SAg epitopes targeted with vaccines are unlikely to bind and provide broad protection against all *S. aureus* superantigens. This may exacerbate issues of non-protective immune imprinting, which has recently shown to prevent the development of protective immunity and result in adverse infection outcomes (*15, 16*). These insights underscore the importance of epitope-specific targeting for effective protection in future vaccines and immunotherapies.

As a proof-of-concept, we engineered multifunctional Nb constructs capable of simultaneously neutralizing SEB, SEC, TSST-1, and Hla. Because clinically encountered S. aureus strains encode heterogeneous combinations of virulence factors, multivalent strategies are thought to provide more reliable functional coverage (104). The aerosolizable trimeric construct C112-C107-H204 demonstrated potent biparatopic SEC neutralization and robust Hla neutralization both in vitro and in vivo. Building on this platform, our decameric construct achieved marked neutralization (e.g., picomolar or higher potencies) across all four targets, establishing the feasibility of comprehensive yet precise multispecific anti-virulence targeting within a single molecular framework. Given its half-life extended Fc YTE scaffold, this construct could enable infrequent dosing for prophylactic use in highly vulnerable patients to prevent or treat bacteremia and pneumonia. Robust *in vitro* neutralizing activities for these 4 toxins were achieved with a single monotherapy construct, which may allow for a more comprehensive anti-virulence treatment against synergistic toxins using a single dose. Multifunctionality may also enhance neutralizing breadth against a wider range of strains, as SAgs are heterogeneously encoded and expressed in different combinations in different *S. aureus* strains (*86, 105–117*). This multivalent Nb Fc fusion platform can be reconfigured or augmented with Nbs against other virulence factors to further tailor or expand therapeutic function, effectively addressing the complex virulence profiles of *S. aureus*.

Therapeutic development against SAgs has been hindered by the lack of predictive animal models. Mice are not native hosts of *S. aureus* and SAgs do not interact strongly with murine MHCII and TCR subtypes, requiring humanized transgenic mouse models (*118*). WT mouse models, which are often used to evaluate Hla driven disease (*39, 40, 103, 119–121*), may underrepresent contributions of SAgs. Rabbit and ferret models are more sensitive but require complex regulatory review and substantially higher per-study costs (*29, 69, 71, 85, 122, 123*). More predictive nonhuman primates remain impractical due to expense and ethical constraints. *In vivo* toxin expression also varies by strain, site, and disease stage(*124*), further complicating model interpretation. Our neutralization panel targeted major clinically important toxins, but given the broad variability in toxin carriage and expression across *S. aureus* lineages, expanding evaluation to diverse clinical isolates will be essential. Closing this persistent model and strain gap will be critical to advancing *S. aureus* therapeutic development.

Together, these findings establish multivalent Nbs as a platform for systematically targeting complex *S. aureus* virulence programs. The high modularity of the Nb format enables rapid incorporation of binders against additional virulence factors, including bicomponent leukocidins and non-classical superantigens, providing a flexible framework for expanding functional coverage. Multivalent Nbs can also be specifically tailored against virulence factors that are variably expressed in specific infection subtypes and timelines, optimizing therapeutic activity against specific diseases. More broadly, this precision immunotherapy strategy could be adapted to other bacterial pathogens with complex, variable virulence profiles, offering a flexible and cost-effective platform to address unmet needs in infectious disease treatment.

## Acknowledgments

We thank Shi lab members, as well as Aneel Aggarwal, Harm Van bakel, and Jeremiah Faith (Mount Sinai) for the insightful discussion, Wilmara Salgado-Pabon (University of Wisconsin), Patrick Schlievert (University of Iowa), James Cassat (Vanderbilt University), Richard Novick (NYU), and Matthew Culyba (University of Pittsburgh) for their expertise. Y.J.K. and Y.S. are inventors on a patent application covering several nanobodies evaluated in this study. All other authors declare no competing interests. Funding: This work is partially supported by a school seed fund (to Y.S.), a T32 training grant (to Y.J.K; 5T32CA078207-24, 5T32GM062754-23), and Project Grant PJT-198030 from the Canadian Institutes of Health Research (to J.K.M.). N.R.B was supported in part by an R.G.E. Murray Graduate Scholarship. Funding for Hook Lab: NIH grant R01HL164821, Cystic Fibrosis Foundation Research Grant 004792G222, and American Lung Association COVID-19 and Emerging Respiratory Viruses Research Award 1031520 all to J.L.H. The authors would like to acknowledge the following NIH S10 Shared Instrumentation grant S10OD032437-01 for supporting this work. The content is solely the responsibility of the authors and does not necessarily represent the official views of the National Institutes of Health.

## Author contributions

Y.J.K and Y.S. conceived the work and drafted the manuscript with input from all authors. Y.J.L performed biochemistry, proteomics and neutralization experiments, with the help of Y.X., Z.S., and M.L. N.R.W. performed the in vivo experiments related to SAg. W.H. performed cryoEM analysis. C.S. and S.M. evaluated Nbs in the Hla murine model. Y.S., J.K.M., J.L.H., K.C., and D.J.T. supervised the study.

## Methods

### Recombinant Toxoid Purification and Immunization

A llama (Capralogics; Hardwick, MA) was immunized using a cocktail of purified recombinant toxoids of Hla, SEB, SEC, and TSST-1. Vaccine constructs of each superantigen included TSST-1 G31R H135A (*71, 73*), SEB L45R Y89A Y94A (*73, 74*), and SEC-2 N23A Y94A (*73, 75*). These toxoids were developed as safe vaccine candidates by abrogating MHCII and TCR binding activity, displaying no *in vitro* or *in vivo* activities (*71–75*). Inactivated Hla toxoid contained the mutation H35L, which prevents the oligomerization of Hla into cytotoxic pores (*68–70*). WT sequences for SEB (residues 28-266, UNIPROT: P01552), SEC-2 (residues 28-266, UNIPROT: P34071), TSST-1 (residues 41-234, UNIPROT: P0A0L2), and Hla (residues 27-319, UNIPROT: P09616) were codon optimized and synthesized into pET19b vectors.

Codon optimized sequences for toxoids were synthesized into pET19b vectors, which contain N-terminal His tags, using BamHI and NdeI restriction sites (Synbio Technologies) and transformed into *E. coli* BL-21 expression systems. Single colonies were incubated overnight in 37°C at 240 rpm in autoclaved LB miller broth media with 50µg/ml of ampicillin salt. Overnight seeds were incubated with large volume LB broth cultures at 37°C to an OD600 of 0.6-0.8 and was induced with 500µM of Isopropyl β-D-1-thiogalactopyranoside (IPTG) for recombinant protein expression. Recombinant protein expression ensued at 16°C at 240 rpm for 16-18 hours. Expression cultures were pelleted, resuspended in lysis buffer (1x PBS, 10mM Imidazole, 0.02% Triton X-100, pH 7.4), and lysed with untrasonication. Lysed cells were clarified through high-speed centrifugation and transferred to pre-washed His-Cobalt resin for 1 hour incubation at 4°C using a rotating mixer. His-cobalt resin was collected onto a filtered column and washed with His-cobalt equilibration buffer (1xPBS, 10mM Imidazole, pH 7.4) until minimal A280nm signal was measured. Recombinant proteins were eluted using ∼5 column volumes of a high imidazole buffer (1xPBS, 150mM Imidazole, pH 7.4). Recombinant toxoids were further purified using size exclusion chromatography (Superdex 75 10/300 GL column connected to AKTA Pure system). Endotoxins were removed from purified toxoids using an immobilized polymyxin B ligand-based resin (Detoxi-Gel Endotoxin Removing Gel; Thermo Fisher).

1mg of each toxoid was administered for the initial immunization bolus in Complete Freund’s Adjuvant, followed by 3 additional boosters of 0.5mg of each toxoids in incomplete Freund’s Adjuvant every 3 weeks. Once a sufficient antibody titer enhancement was observed (∼2 log enhanced titers) in a test bleed, whole blood was requested for antibody pull-down experiments and PBMC cDNA next generation sequencing (NGS) library construction.

### Whole Blood Processing for PBMC and Plasma Isolation

Around 500ml of anticoagulated immunized whole blood is received and processed immediately for 1) proteomic analysis of antibody pull-down experiments and 2) generation of cDNA genomic libraries of circulating PBMC’s. Whole blood was diluted 1:3 (v/v) with cell culture grade Dulbeco’s PBS (dPBS) and filtered through a 70µm mesh cell sieve to remove any blood clots. 15 ml of Histopaque density gradient media was added to each 50ml Falcon tube, and 30-35 ml of the 1:3 diluted whole blood was gently layered on top of the Histopaque media. Samples were centrifuged at 1000xg for 25 minutes at room temperature with gentle acceleration and with no brakes engaged. The diluted plasma was carefully isolated into 50ml Falcon tubes and kept at 4°C for antibody isolation and antigen affinity pulldown experiments. Separated PBMCs on top of the Histopaque media were isolated using a plastic dropper and transferred to a 50ml Falcon tube on ice, minimizing the pickup of excess Histopaque media. Isolated PBMCs were washed and pelleted (200xg for 5 minutes) twice with dPBS for mRNA extraction and cDNA library generation.

### PBMC cDNA library preparation

Around 1×10^9^ PBMCs were used to extract mRNA using RNeasy kit (Qiagen) and reverse-transcribed into cDNA using Maxima H Minus cDNA Synthesis Master Mix (Thermo). In the first PCR reaction to amplify the VH domain of camelid IgG’s, the forward primer CALL001 (5’-GTCCTGGCTGCTCTTCTACAAGG-3’; (*125*)) hybridizes with a conserved leader signal for VH subgroup III whereas the reverse primer CH2FORTA4 (5′-CGCCATCAAGGTACCAGTTGA-3′; (*126*)) hybridizes with the CH2 region of camelid IgG mRNA. The camelid IgG1 populations that are amplified include 1) the VH and hinge regions of hcAbs and 2) the VH, CH1, and hinge regions of traditional IgG1 antibodies. The amplicons were separated with gel electrophoresis using a 1% agarose ethidium bromide gel, and the lower molecular weight amplicon corresponding to the VH region of hcAbs were isolated using gel extraction (Monarch NEB). In the 2^nd^ PCR reaction, just the VH domain of hcAbs was further isolated starting with a forward primer for FR1 (5’-ATCTACACTCTTTCCCTACACGACGCTCTTCCGATCTNNNNNNNNATGGCT[C/G]A[G/T]GTGCAGCTGGTG

GAGTCTGG-3’) and a reverse primer for FR4 (5’-ATCTACACTCTTTCCCTACACGACGCTCTTCCGATCTNNNNNNNNATGGCT[C/G]A[G/T]GTGCAGCTGGTG

GAGTCTGG-3’). In the 3rd PCR reaction, P5 (forward primer P5 MiSeq: 5’-AATGATACGGCGACCACCGAGATCTACACTCTTTCCCTA-3’) and P7 (reverse primer P7 MiSeq: 5’-CAAGCAGAAGACGGCATACGAGATTTCTGAATGTGACTGGAGTTCA-3’) Illumina adapter sequences were added to aid Illumina MiSeq cluster identification. The final amplicons were sequenced using an Illumina Miseq platform with 300bp paired-end model.

### Heavy-Chain-Only Antibody Isolation

The plasma isolated from the immunized whole blood is converted to sera by clearing clotting factors with the addition of a final concentration of 10mM calcium chloride. Clotting factors are allowed to precipitate through incubation at room temperature for 1-2 hours. Precipitated clotting factors are removed by centrifuging at 4300xg for 30 minutes. The isolated sera are diluted with dPBS to 10x dilution of the original whole blood and was mixed with Protein G resin (10% of the diluted serum volume in resin bed volume). Serum was allowed to incubate with Protein G resin at 4°C using a rotating mixer for 1 hour. Protein G resin was loaded on a filtered column and washed with dPBS until minimal A280 signal was observed. Heavy-chain-only antibodies were eluted using a gentle sodium acetate elution buffer (0.1M Sodium Acetate, 0.5M NaCl, pH3.0), followed by the elution of traditional IgG1 antibodies using a harsher glycine-based elution buffer (0.1M glycine-HCl, pH 3.0). The flow through containing additional heavy-chain-only antibodies was incubated with Protein A resin for 4°C using a rotating mixer for 1 hour, washed with dPBS, and eluted with glycine-based elution buffer to isolate remaining hcAbs. Antibodies eluted with sodium acetate elution buffer were neutralized to pH 7.4 using 1M Tris, whereas antibodies eluted with glycine elution buffer were neutralized to pH 7.4 using 1M sodium hydroxide. Purified hcAbs were pooled and dialyzed at 4°C overnight using a 10 kDa dialysis cassette in dPBS. Pooled hcAbs were adjusted to a concentration of 1 mg/ml for antibody pulldown experiments.

### Antigen Specific hcAb Pulldown

Wild type SEB, SEC, TSST-1, and Hla were synthesized in pET19b vectors and produced as previously described (*53*) and dialyzed into Cyanogen Bromide (CnBr) resin coupling buffer (100mM NaHCO_3_, 500mM NaCl, pH 8.3). 300ul of CnBr resin was prepared according to manufacturer protocols and incubated with 1.5-4 mg of recombinant toxin for free amine conjugation at 4°C overnight using a rotating mixer. Antigen immobilized CnBr agarose resin was blocked and prepared according to manufacturer protocols. 40-60mgs of the isolated hcAbs were incubated with the antigen immobilized CnBr resin using a rotating mixer at 4°C for 1 hour. The antigen immobilized resin was collected in a filtered column and washed with a high salt PBS buffer (1xPBS, 0.5M NaCl). Isolated heavy-chain-only antibodies were eluted stepwise with increasing concentrations of magnesium chloride (1, 2, 3, and 4.5M). Starting with 1M magnesium chloride, the resin was incubated with 3x volume of each elution condition for 5 minutes on a rotating mixer at room temperature and eluted into a 1.5ml Eppendorf tube through 50xg centrifugation for 30 seconds. Eluted antibodies were buffer exchanged into 0.2x PBS using a conical 10 kDa concentrator. Protein concentrations were measured for each fraction and flash frozen in liquid nitrogen for downstream mass spectrometry sample preparation.

### Mass Spectrometry Sample Preparation and Analysis

Flash frozen eluted antibody samples were lyophilized using a SpeedVac and reconstituted in a denaturing solution (8M urea, 5mM DTT, 5mM TCEP). After heating and dilution, the antibodies were digested by either trypsin or chymotrypsin, desalted, and analyzed by LC-MS. Purified peptides were analyzed with a Vanquish NEO coupled with a Q Exactive Exploris 480 Orbitrap mass spectrometer (Thermo Fisher). Peptides were loaded onto an analytical column and eluted using a 60-min liquid chromatography gradient (3% B–5% B, 0–2 min; 5% B–25% B, 2–47 min; 25% B–80% B, 47 – 54 min; 80% B - 3% B, 54 min - 54 min 10 sec; 3% B, 54 min 10 sec - 60 min; mobile phase A consisted of 0.1% formic acid (FA), and mobile phase B consisted of 0.1% FA in 100% acetonitrile (ACN)). CDR peptides and the corresponding Nb sequences were identified and quantified by AugarLlama to deconvolute specific Nb sequences, as described (*53*).

### Recombinant Nb and Recombinant Toxin Purification

Sequences for identified Nbs were codon optimized for *E. coli* ClearColi BL-21 expression systems and synthesized into a pET-21b vector using restriction enzymes EcoRI and HindIII (Synbio Technologies). Recombinant nanobodies were expressed with an N-terminal T7 tag and C-terminal His tag. Recombinant toxins were synthesized in pET-19b vectors. Non-Fc fused multivalent Nbs were synthesized into pET21b through a CRO service (Synbio) or subcloned within the lab. For subcloned constructs, Nbs were PCR amplified using primer pairs that add different restriction enzyme sites. Gene fragments of GGGGSx5 linkers with compatible restriction sites and flanking PCR amplification sequences were synthesized (IDT Bioservices) and PCR amplified. All Nb and linker amplicons were digested with appropriate restriction enzymes and ligated using a 5 insert ligation mixture with T4 ligase. Subcloned multivalent Nb constructs were sequence verified using Sanger Sequencing (GeneWiz).

BL-21 seeds were grown in LB broth cultures at 37°C to an OD600 of 0.6-0.8 and was supplemented with 500µM final concentration of Isopropyl β-D-1-thiogalactopyranoside (IPTG) for recombinant protein expression at 16°C for 16-18 hours. Expression cultures were pelleted (4300xg for 15 minutes), resuspended in lysis buffer (1x PBS, 10mM Imidazole, 0.02% Triton X-100, pH 7.4), and ultrasonicated using a wand ultrasonicator. Lysed cells were clarified through high-speed centrifugation (21,000-30,000xg for 10 minutes) and mixed with pre-washed His-Cobalt resin for 1 hour at 4°C using a rotating mixer. His-cobalt resin captured Nbs were washed with His-cobalt equilibration buffer (1xPBS, 10mM Imidazole, pH 7.4) and eluted using ∼5 column volumes of a high imidazole buffer (1xPBS, 150mM Imidazole, pH 7.4). Recombinant Nbs were further purified of endotoxins using an immobilized polymyxin B ligand-based resin (Detoxi-Gel Endotoxin Removing Gel; Thermo Fisher).

### Mammalian Cell Expression of Fc Fusion Nbs

Nanobody Fc fusion constructs were cloned into a proprietary expression plasmid vector with an N-terminal secretion signal peptide and an ampicillin resistance gene for cloning selection. Plasmids were purified from positive clones, verified with Sanger Sequencing (Genewiz), and purified at Midiprep scales for transient transfection.

Nb Fc fusion constructs were expressed in Expi293F cells (Thermo Fisher), a high-yield transient expression system based on engineered HEK cell line. Frozen Expi293 cells were thawed, washed, and cultured in Expi293 expression media in a 37°C incubator with 8% CO2. Cells were grown to ∼3-5×10^6^ cells/ml and split to 0.5×10^6^ cells/ml until the desired cell density and volume were achieved. Expi293F cells were diluted to 2.9×10^6^ cells/ml and transiently transfected with Nb-Fc plasmids pre-incubated with Expifectamine 293 reagent in Opti-MEM reduced serum media. Transiently transfected Expi293F cells were incubated with a shaker at 180 rpm in a 37°C incubator with 8% CO2 for 20 hours, at which time the Expi293F culture was enhanced using ExpiFectamine 293 enhancer 1 and ExpiFectamine 293 enhancer 2. Expi293F cells were allowed to express proteins on a shaker at 180 rpm in a 37°C incubator with 8% CO2 for 4-6 days.

Expression cultures were transferred to 50ml falcon tubes and pelleted at 300xg for 3 minutes. Supernatants were isolated and mixed with recombinant Protein A agarose resin using a tumbling incubator for 1 hour at 4°C. Protein A resin captured Nb Fc fusion proteins were transferred to a filtered column and washed with 1xPBS. Washed Nb Fc fusion proteins were eluted with 5 resin bed volumes of 0.1 M glycine pH 3.0, neutralizing the pH with 1M Tris pH 8.0. Eluted Nb Fc fusion constructs were buffer exchanged into cell culture grade dPBS using a 30kDa molecular weight cut off concentrator. Endotoxin levels were verified using a LAL based assay (ToxinSensor Chromogenic LAL Endotoxin Assay kit, Genscript) or Recombinase C based assay (Recombinant Factor C Endotoxin Detection Kit, ACROBiosystems). Detoxi-Gel endotoxin removing gel was used to remove any excess endotoxin in Nb Fc fusion proteins before animal studies.

### ELISA Serum Titer and Nb Binding Measurement

Serum titer experiments were conducted with a sandwich ELISA based design. 96-well ELISA plates were coated with 1-10 µg/ml of antigens of interest in a coating buffer (15mM Na_2_CO_3_, 35mM NaHCO_3_, pH 9.6) at 4°C overnight. Non-immobilized antigens were decanted, and the plate was washed with PBST buffer (1x PBS, 0.05% Tween v/v). Plates were blocked (PBST, 5% milk powder w/v) at room temperature for 2 hours, washed, and incubated with serial dilutions of T7-tagged Nbs or serum in blocking buffer at room temperature for 2 hours. Immobilized Nb-antigen complexes were washed and incubated at room temperature for 1 hour with 1:10,000 dilution of HRP-conjugated polyclonal anti-llama IgG1 secondary antibody (for serum titer measurements). Plates were washed thoroughly with PBST and incubated with TMB substrate (R&D Systems) for 10-15 minutes and quenched with STOP solution (R&D Systems). HRP substrate colorimetric signals were measured using a plate reader at wavelength 450 nm, using 550 nm as a background reference wavelength. ELISA curves were plotted on GraphPad Prism and analyzed using a nonlinear regression 4PL regression analysis.

Nanobody binding and serum titer experiments were conducted with a sandwich ELISA based design. 96-well ELISA plates were coated with 1-10 µg/ml of antigens of interest in a coating buffer (15mM Na_2_CO_3_, 35mM NaHCO_3_, pH 9.6) at 4°C overnight. Non-immobilized antigens were decanted, and the plate was washed with PBST buffer (1x PBS, 0.05% Tween v/v). Plates were blocked (PBST, 5% milk powder w/v) at room temperature for 2 hours, washed, and incubated with serial dilutions of T7-tagged Nbs or serum in blocking buffer at room temperature for 2 hours. Immobilized Nb-antigen complexes were washed and incubated at room temperature for 1 hour with 1:7500 dilution of HRP-conjugated polyclonal anti-T7 secondary antibody (for Nb binding experiments) or 1:20,000 dilution of HRP-conjugated polyclonal anti-human IgG1 Fc secondary antibody. Plates were washed thoroughly with PBST and incubated with TMB substrate (R&D Systems) for 10-15 minutes and quenched with STOP solution (R&D Systems). HRP substrate colorimetric signals were measured using a plate reader at wavelength 450 nm, using 550 nm as a background reference wavelength. ELISA curves were plotted on GraphPad Prism and analyzed using a nonlinear regression 4PL regression analysis.

### Surface Plasmon Resonance

All SPR measurements were conducted on a Biacore 3000 SPR system (GE Healthcare). Recombinant His tagged SEC in a 10mM sodium acetate buffer (pH 4.0) was immobilized using amine coupling chemistry on a CM5 sensor-chip (Cytiva) activated with EDC/NHS. Excess reactive groups were deactivated with ethanolamine (1M ethanolamine-HCl pH 8.5). Each Nb was prepared in different dilutions in HBS-EP running buffer (GE-Healthcare) and injected at a flow rate of 20µL/min for 180 seconds, followed by a dissociation time of 20-25 minutes. Between each replicate injection, the CM5 sensor chip was regenerated with a 10mM glycine-HCl buffer (pH 1.5-2.0) using a flow rate of 30-40µL/min for 30-45 seconds. Measurements for each concentration were conducted in triplicate. Sensograms were analyzed using BIAevaluation software by using 1:1 Langmuir model fitting or 1:1 Langmuir model fitting with baseline drift correction.

### BioLayer Interference (BLI)

Binding kinetics were measured using the Sartorius Octet R8. Recombinant His-tagged SEC2 was diluted to 10 ng/µL in 10 mM sodium acetate buffer (pH 4.0) and immobilized onto Sartorius AR2G sensors via amine-coupling chemistry activated with EDC/NHS. Residual reactive groups were quenched using 1 M Tris-HCl (pH 7.5) to terminate coupling. Running buffer for all kinetic measurements was PBS. Nanobodies (Nbs) were prepared at a starting concentration of 10 nM and subjected to seven-point 1:2 serial dilutions in PBS to generate the final concentration series. Association was monitored for 300 s, followed by dissociation for 20 mins. New sensors were used for each replicate. All measurements were performed in technical duplicate using independent immobilization cycles, and data were analyzed with 1:1 Langmuir binding model fitting using.

### Recombinant Superantigen PBMC Stimulation Assays

Frozen aliquots of pooled human PBMC cells (Cytologics) were thawed using a 37°C water bath. PBMCs were equilibrated in complete RPMI (RPMI 1640, 10% FBS, 0.3 g/L L-glutamine, 1x Penicillin/Streptomycin) in an 8% CO_2_ incubator at 37°C for 5 minutes, pelleted at 300xg for 3 minutes, and washed with cell culture grade dPBS. PBMCs were further equilibrated in complete RPMI for at least 1 hour in an 8% CO_2_ incubator at 37°C before stimulation with recombinant superantigen. Endotoxin removed Nbs were serially diluted in cRPMI and mixed with a dose response optimized concentration of recombinant SEB, SEC, and TSST-1. Nb-SAg binding mixtures were allowed to incubate at room temperature for 30 minutes. Nb-SAg samples were mixed with 1×10^5^ cells per well in a tissue culture grade 96-well plate at a final SAg concentration of 0.01-0.1nM SEB or SEC and 1-10nM TSST-1. PBMCs were allowed to be stimulated in an 8% CO_2_ incubator at 37°C for 18 hours. Media supernatants containing PBMC secreted cytokines (IFN-γ) were collected for downstream ELISA measurements.

IFN-γ concentrations were measured using a commercial ELISA based IFN-γ measurement kit (ELISA Max Deluxe Set Human IFN-γ; Biolegend). An ELISA 96-well plate was coated 1:200 diluted IFN-γ capture antibody in coating buffer (15mM Na_2_CO_3_, 35mM NaHCO_3_, pH 9.6) at 4°C overnight. Plates were washed with PBST and blocked with an albumin-based blocking buffer (2% w/v bovine serum albumin in PBST) for 1 hour at room temperature. Plates were washed and incubated at room temperature for 2 hours with secreted supernatants diluted in albumin-based blocking buffer (1:6 dilution for SEB or SEC and 1:2 dilution for TSST-1). Plates were washed with PBST and incubated with 1:200 dilution of biotinylated IFN-γ detection antibody. Excess detection antibody was washed with PBST and incubated at room temperature for 30 minutes with 1:1000 dilution of HRP-Avidin. The sandwich ELISA immune complex was thoroughly washed with PBST and allowed to react with TMB substrate (R&D Systems) for 10-15 minutes at room temperature. HRP substrate colorimetric signals were measured using a plate reader at wavelength 450 nm, using 550 nm as a background reference wavelength.

Interpolations of IFN-γ standards were used to convert A450 signals to cytokine concentrations. Secreted cytokine concentrations were plotted against Nb concentration on GraphPad Prism and analyzed using a nonlinear regression 4PL regression analysis to determine the neutralization efficacy (neutralization %) and potency (IC_50_).

Overnight S. aureus supernatant experiments were conducted similarly to recombinant SAg experiments. Overnight stationary phase *S. aureus* cultures (MN8, MN8 TSST1 deletion, COL, COL SEB deletion) were pelleted to isolate supernatants containing secreted exotoxins. Optimized dilutions of supernatants were preincubated with 500nM decameric Nb Fc and administered to human PBMCs as previously described. ELISA based cytokine readout kits were used to measure IL-2 levels from overnight cell culture media supernatants. Multiple comparison analyses were conducted with GraphPad using one way ANOVA analyses.

### Rabbit Blood Hemolysis Experiments

Rabbit Alsever’s solutions (Colorado Serum Company) were pelleted through centrifugation (300xg for 3 minutes) and washed with with 1xPBS. Washed rabbit RBC’s were diluted to a final 5% RBC solution (v/v) with PBS. Recombinant Hla was incubated with serial dilutions of Nbs in 1xPBS at room temperature for 30 minutes using non-specific Nbs as controls. Nb-Hla samples were mixed 1:1 with 5% (v/v) RBC solutions in a 96-well plate and incubated at 37°C for 30 minutes for hemolysis to ensue. Hemolyzed samples were centrifuged to collect hemoglobin-containing supernatants, which were transferred to a 96-well plate for hemoglobin absorbance measurement (542nm). Using blank and nonspecific Nb controls, absorbance values were converted to percent hemolysis and neutralization values, which were plotted on Graphpad Prism. Neutralizing IC50’s were calculated usin nonlinear regression 4PL analysis.

### Size Exclusion Chromatography Epitope Binning

Size exclusion chromatography experiments were done on a Superdex 75 10/300 GL increase column connected to an AKTA Pure FPLC system as well as on a Shodex Protein LW-803 column connected to a Shimadzu HPLC system. All possible 2 Nb combinations were incubated with antigens of interest in 1xPBS for at least 30 minutes at room temperature before size exclusion chromatography experiments. Elution profiles for two Nb-antigen complexes were compared to elution profiles for one Nb-antigen complexes and antigen only controls.

### Cryo-EM Structure Sample preparation

Purified recombinant Nb-antigen co-complexes were used for structural determination by Cryo-EM. Two Nb occupying different epitopes were allowed to incubate with antigens each at a 2:1 molar ratio. Nb-antigen co-complexes were allowed to form at room temperature in 1xPBS for at least 30 minutes to ensure sample homogeneity. Antigen-Nb co-complexes were purified with size exclusion chromatography. Ultrafoil R1.2/1.3 Au 300 mesh grids (Quantifoil) were negatively glow discharged for 30 seconds at 15 mA. ThermoFisher Vitrobot Mark IV was set to 4°C and 100% humidity during freezing. The sample was diluted to 0.6 mg/mL; 3.5 µL of sample was applied to the grid and blotted for 2 seconds with a blot force of 0.

### CryoEM data collection of SEC-Nbs complex

The frozen-dehydrated grids were transferred to a Titan Krios (Thermo Fisher Scientific) transmission electron microscope, equipped with a Gatan K3 direct-electron counting camera and a BioQuantum energy filter, for data acquisition under strict light-controlled conditions. Specimen movies were recorded using SerialEM with a beam-tilt image-shift data collection strategy in a 3 × 3 pattern, capturing one shot per hole at a nominal defocus range of −0.8 to −2.0 μm. The K3 camera operated in super-resolution mode at a nominal magnification of 81,000×, yielding a physical pixel size of 1.07 Å/pixel. Each movie stack consisted of 50 frames, with a total exposure time of 2.5 seconds (0.05 seconds per frame). The accumulated electron dose was approximately 44 electrons/Å² per stack.

### CryoEM Image processing of SEC-Nbs complex

Movie stacks of the SEC-Nbs complex were processed using CryoSPARC (version 4.6.0) (Punjani et al., 2017, 2020). Patch motion correction with an F-crop factor of 0.5 was used to align the movie stacks, and patch CTF estimated the contrast-transfer function (CTF) parameters. Particles were autopicked using a 100 Å Gaussian blob. The number of bin2 particles selected after 2D classification is detailed in Extended Data Table 1. Initial 3D volumes and decoys were generated via ab initio reconstruction using selected good 2D classes and unselected bad 2D classes (class number k=3), respectively. The particles cleaned up from 2D classification underwent one round of heterogeneous refinement using the ab initio 3D volume from good 2D classes and decoy 3D volumes from bad 2D classes. Bin1 particles of SEC-Nbs were then re-extracted from dose-weighted micrographs based on their coordinates and angular information. The final particle set underwent non-uniform 3D refinement, followed by global and local CTF refinements, and local 3D refinement, resulting in final maps with global resolutions determined using the 0.143 gold-standard Fourier shell correlation (FSC) criterion (Extended Data Table 1). Half maps were used to determine the local resolution of each map using Relion 5.0 (Fernandez-Leiro & Scheres, 2017; Zivanov et al., 2018). The final 2.7 Å cryoEM map was then used for model building and refinement.

### Model building and refinement

The AlphaFold 3 predicted SEC-Nbs complex model was docked into cryo-EM map using Chimerax (Pettersen et al., 2021). The final model was manually built in coot (Emsley & Crispin, 2018) and subjected to real-space all-atom refinement in Phenix (Afonine et al., 2018). The final refinement statistics for the models are provided in Extended Data Table 1 (*127–133*).

### AlphaFold 3 Modeling

Amino acid sequences for Nbs and Hla were inputted together into AlphaFold 3 servers to generate structural models. Nb-antigen models with iPTM (interface Predicted TM-score) score thresholds of 0.8 or above were considered high confidence models. CIF files of AlphaFold 3 models were displayed on ChimeraX for downstream analyses.

### Transgenic Humanized HLA-DR4 Mouse Models of Bacteremia and Sepsis

Humanized DR4-B6 transgenic mice were assessed for SAg neutralization, as SAgs do not interact strongly with endogenous mouse MHCII receptors. Transgenic male and female HLA-DR4-IE (DRB1*0401) humanized C57BL/6 (B6) mice were used for the *in vivo* experiments. The prototypical clinical strain SEC (MW2) were used for these models, which have been thoroughly used in our previous work (*84, 89, 134, 135*). 1×10^6^ CFUs of *S. aureus* strain MW2 (SEC+) was injected through tail vein injection. Mice were sacrificed at 72 hours post infection (hpi) and bacteria were enumerated by plating liver and kidney homogenates on mannitol salt agar (*84*). Using at least 10 animals per arm, therapeutic Nbs were compared against non-specific Nb controls to assess protection against *S. aureus* bacteremia. To assess efficacy in Nbs unoptimized for half-life, Nbs were administered at a dose of 10 mg/kg intraperitoneally 4 hours prior and 6 hours after intravenous *S. aureus* inoculation. Weight loss was monitored at 24 intervals for 3 days. All animals were euthanized for CFU enumeration in the liver and kidney. These experiments were in accordance with the Canadian Council on Animal Care Guide to the Care and Use of Experimental Animals, and the animal protocol (AUP-2024-082) was approved by the Animal Use Subcommittee at the University of Western Ontario.

### WT Swiss Webster Models of Bacterial Pneumonia

#### Animals

Mice were male Swiss Webster, purchased from Charles River Laboratories, 30-45 g, and 4-8 weeks old.

#### Bacterial strain and preparation

*S. aureus* was GFP-expressing USA300 LAC (SA). SA were stored at -80°C in 25% glycerol in autoclaved Luria-Bertani (LB) broth media (MP Biomedicals) and propagated on LB-agar plates containing chloramphenicol (10 ug/mL). Plates were refreshed from frozen stock every 2 weeks. Single SA colonies were propagated in autoclaved LB media containing chloramphenicol (10 μg/mL) in a shaking incubator at 37°C and 200 rpm (New Brunswick Scientific) for 18 h (stationary growth phase). SA was prepared for intranasal instillation by centrifuging 650 μL of the stationary growth phase culture, then resuspending it in 150 μL of Nb-containing DPBS.

#### Intranasal instillation

SA instilled within 30 min of removing the bacteria from the incubator. Mice were weighed, then anesthetized with inhaled isoflurane (4%) and intraperitoneal injections of ketamine (up to 1 mg) and xylazine (up to 0.1 mg). Each mouse was instilled with 30 μL of prepared Nb-SA solution to deliver 1 x 10^8^ CFU per mouse. Instillation quality was recorded at the time of instillation by the performing investigator and considered acceptable for experiments if no loss of instillate was observed. Mice woke from anesthesia within 3 min of instillation.

#### Breathing score

A blinded investigator assessed and recorded mouse breathing score on an hourly basis for 3 h after Nb-SA instillation. No mice met our IACUC-approved criteria for need for euthanasia during the experimental procedures. All mice were euthanized at the conclusion of the experiments.

#### Lung wet weight to body weight (LW/BW) ratio and lung bacterial content quantifications

LW/BW ratio and lung bacterial content were quantified in the same experiment. At 3 h after SA instillation, we used our established methods (*97*) to exsanguinate anesthetized mice by cardiac puncture, then excise and weigh the lungs. The lungs were mechanically homogenized by crushing in a specimen bag, then diluted in 1 mL of DPBS containing Ca^2+^ and Mg^2+^. We quantified SA CFU by serial dilutions on chloramphenicol-containing LB agar plates.

#### Statistics

Statistics are indicated in figures and legends. Paired comparisons were analyzed using two-tailed *t* tests. We considered statistical significance at *p* < 0.05. Data were analyzed and figures were prepared using Microsoft Excel and PRISM (GraphPad, version 10.5.0).

#### Study approval

The Institutional Animal Care and Use Committee of the Icahn School of Medicine at Mount Sinai approved the procedures related to mouse models of lung infection.

