## Supplemental Figures for "Multivalent Nanobodies for Potent and Broad Neutralization of *Staphylococcus aureus* Toxins"

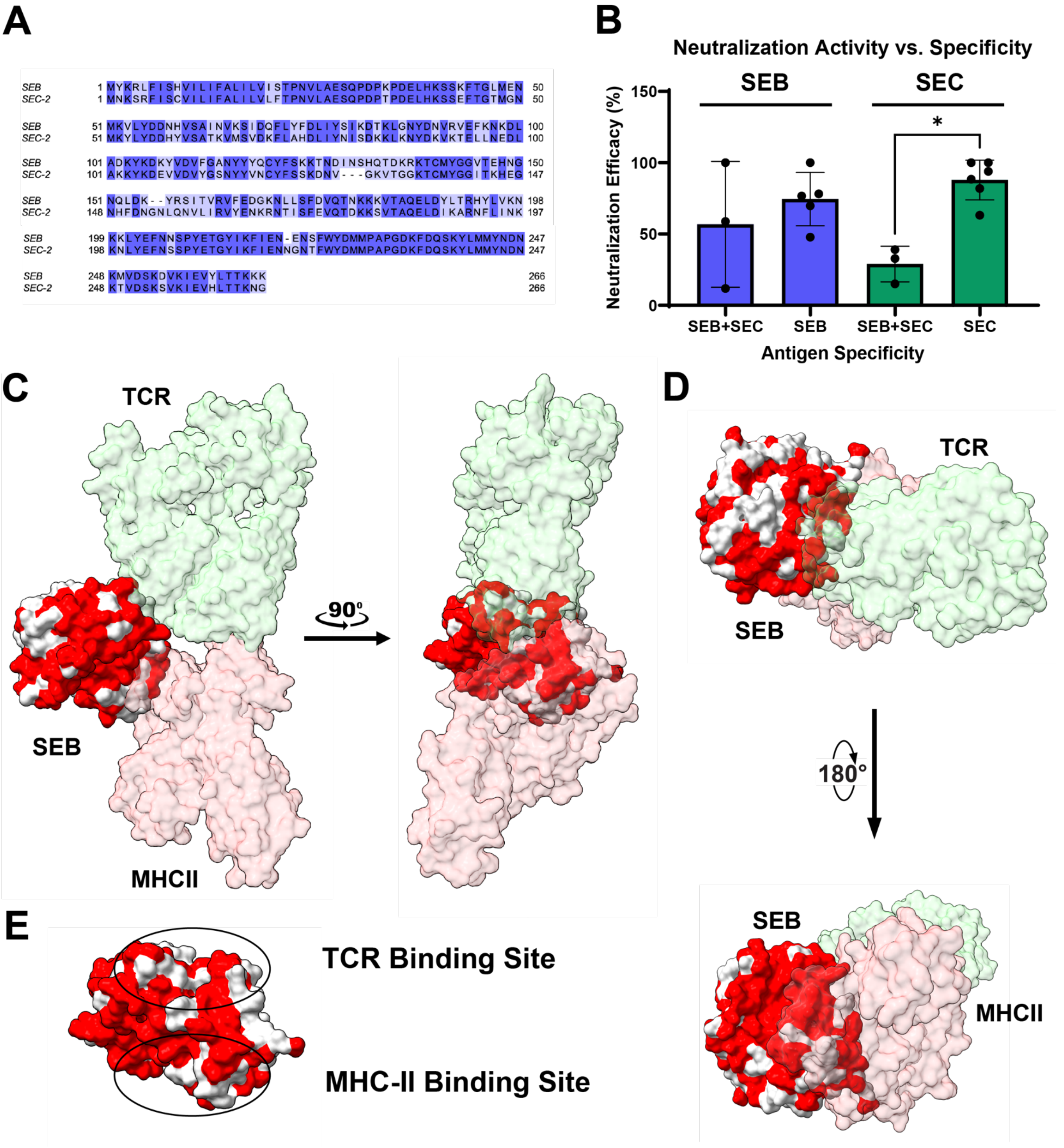


**Supplemental Figure 1.** (**A**) Sequence alignment of SEB and SEC by Clustal Omega. Overall sequence homology is 66.5%. (**B**) Neutralizing efficacies of Nbs cross-reactive to SEB and SEC compared to neutralizing efficacies of monospecific Nbs (**C-D**) Structure of SEB in complex with TCR and MHCII (PDB: 4C56) with residues shared by SEB and SEC highlighted in red. TCR and MHCII interfacing epitopes are less contiguously conserved between SEB and SEC. (**E**) TCR and MHCII sites highlighted showing conserved (red) and non-conserved (grey) residues between SEB and SEC.

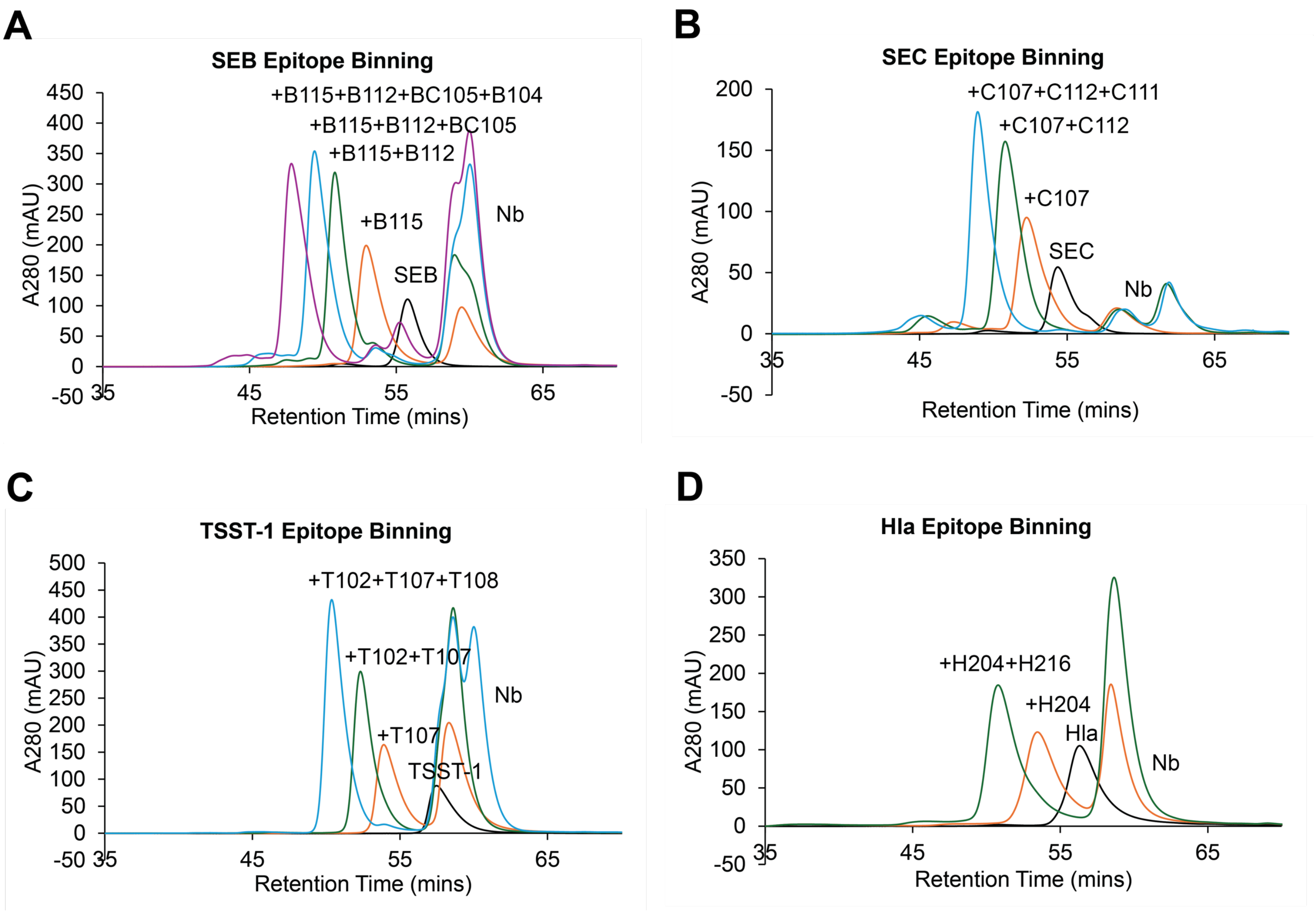


**Supplementary Figure 2.** Representative size exclusion chromatography chromatograms for nanobodies against **(A)** SEB (4 epitopes), (**B**) SEC (3 epitopes) (**C**) TSST-1 (3 epitopes) and (**D**) Hla (2 epitopes).

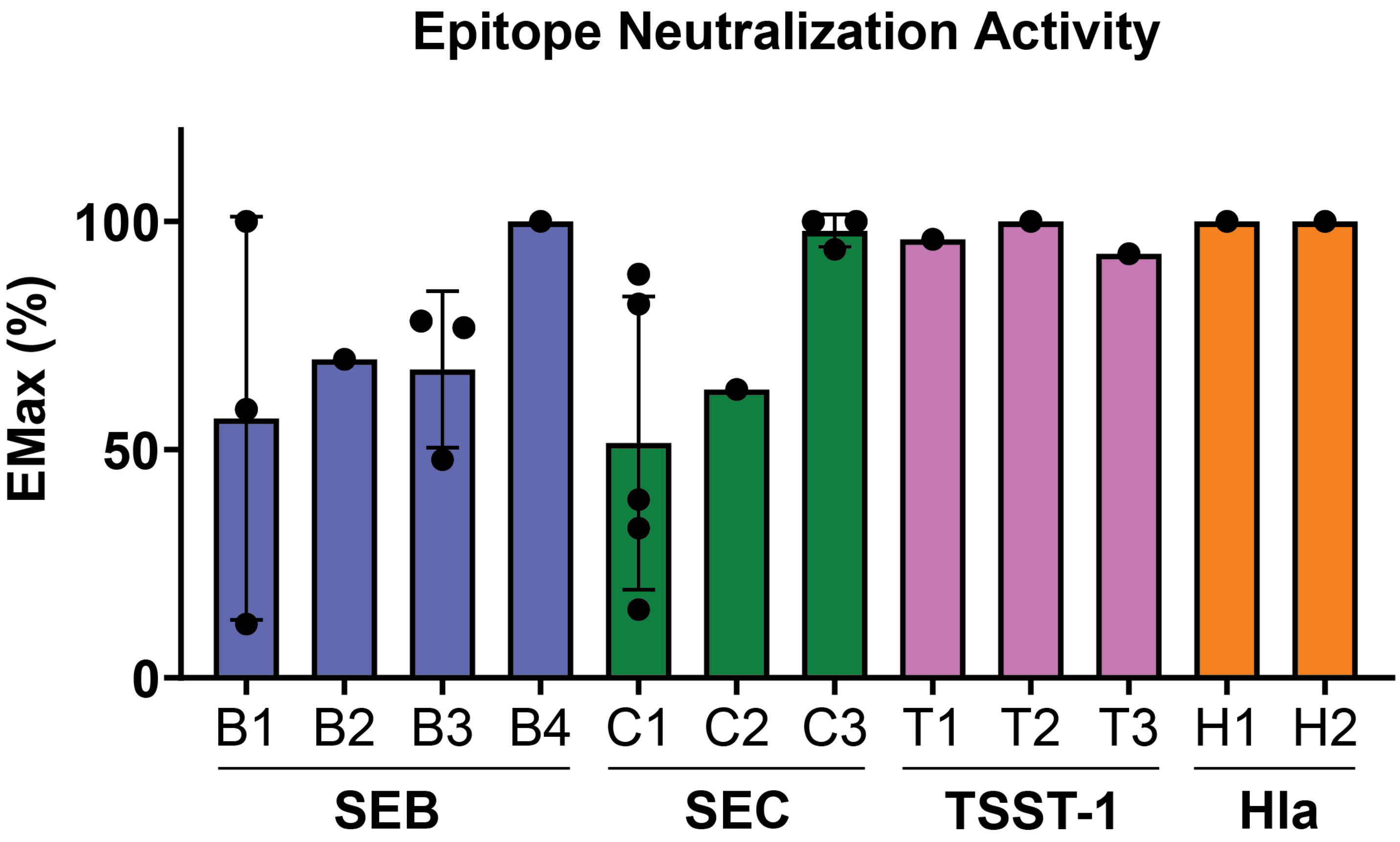


**Supplementary Figure 3.** Neutralization Emax (%) plotted according to toxin and epitope class.

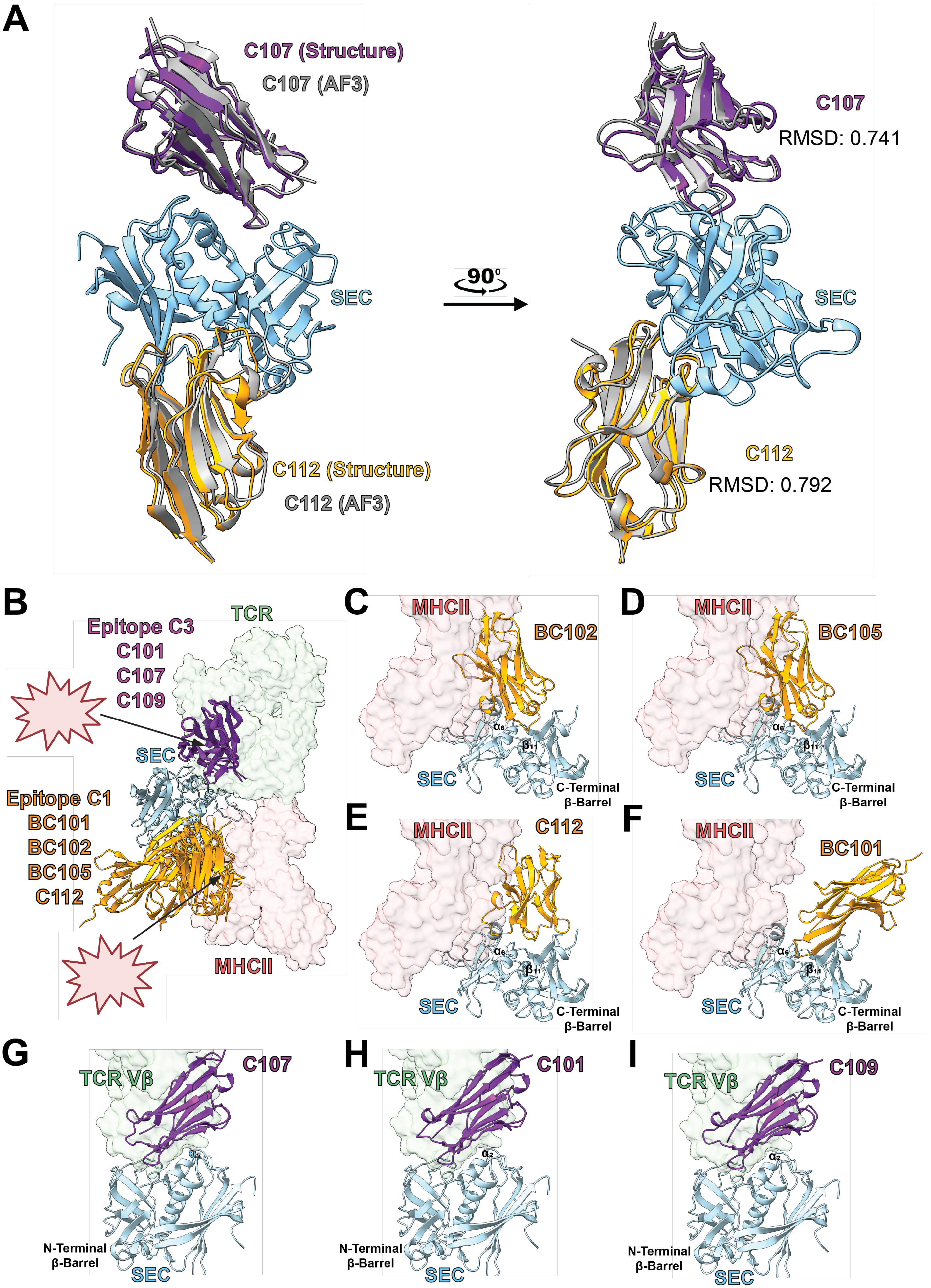
A
C107 (Structure)
C107 (AF3)
SEC
C112 (Structure)
C112 (AF3)
90°
C107
RMSD: 0.741
SEC
C112
RMSD: 0.792
B
TCR
Epitope C3
C101
C107
C109
SEC
Epitope C1
BC101
BC102
BC105
C112
MHCII
C
MHCII
BC102
SEC
C-Terminal
β-Barrel
D
MHCII
BC105
SEC
C-Terminal
β-Barrel
E
MHCII
C112
SEC
C-Terminal
β-Barrel
F
MHCII
BC101
SEC
C-Terminal
β-Barrel
G
TCR Vβ
C107
N-Terminal
β-Barrel
SEC
H
TCR Vβ
C101
N-Terminal
β-Barrel
SEC
I
TCR Vβ
C109
N-Terminal
β-Barrel
SEC

**Supplementary Figure 4. AlphaFold 3 (AF3) models of anti-SEC Nbs.** (**A**) Superimposition of Cryo-EM structures and high confidence AF3 models of SEC in complex with C107 and C112. **(B**) All high confidence AF3 models of SEC Nbs (iPTM>0.8) overlayed on SEC-2. SAg-TCR structures were superimposed from PDB codes 4C56 and 1JCK. SAg-MHCII structures were superimposed from PDB code 1JWM (**C-F**) AF3 models of epitope C1 Nbs, which interact with beta strand 11 and alpha helix 6 to block MHCII binding (**G-I**) AF3 models of epitope C3 Nbs, which interact with alpha helix 2 and the N-terminal beta barrel domain to block TCR binding.

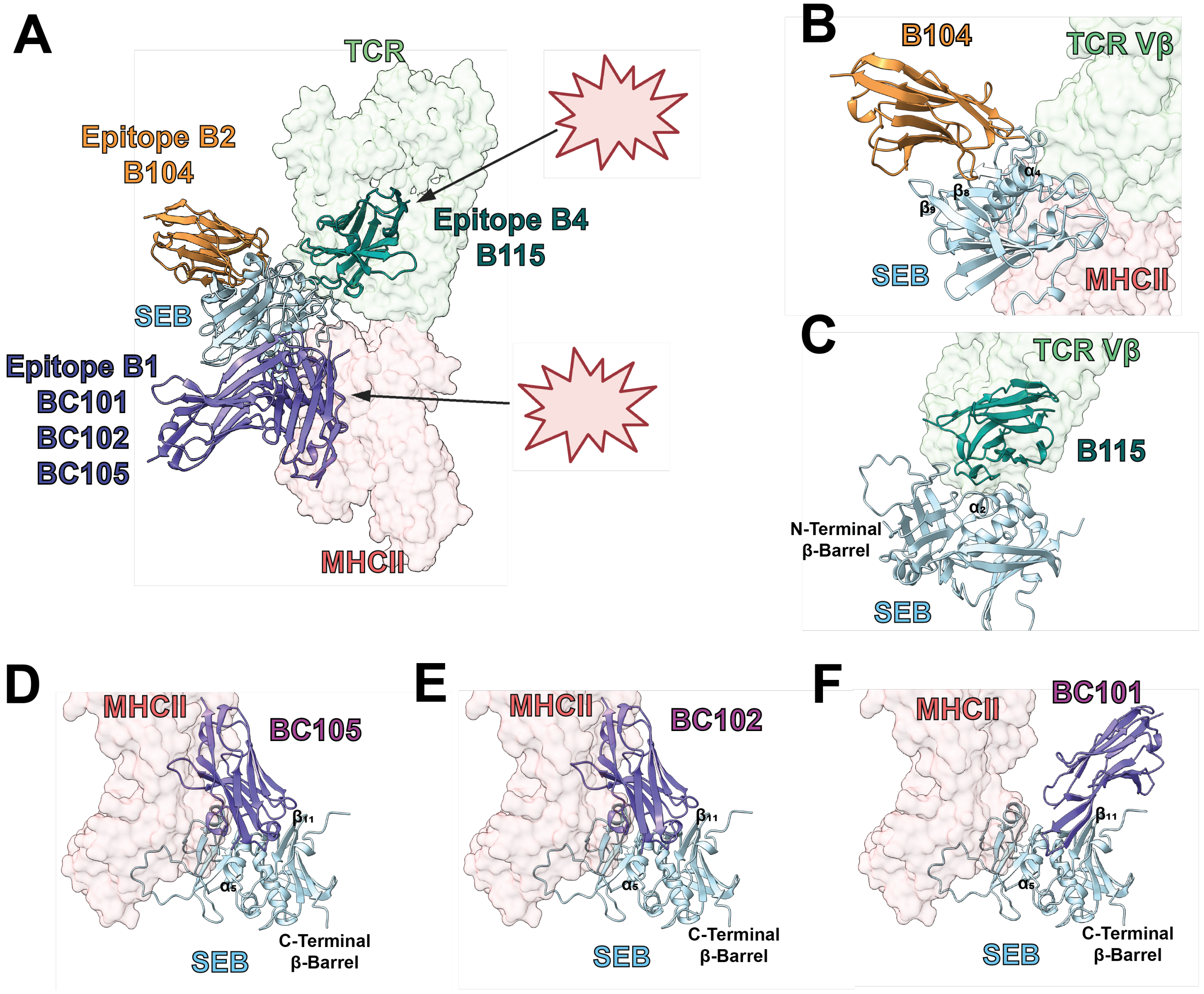


**Supplementary Figure 5. AlphaFold 3 (AF3) models of anti-SEB Nbs (A)** All high confidence AF3 models of SEB Nbs (iPTM>0.8) overlayed on SEB. SAg cocomplex structures with TCR and MHCII were superimposed from PDB code 4C56 (**B**) AF3 models of epitope B2 Nbs B104 which interact with beta strand 8 and 9 and alpha helix 4. B104 does not block TCR/MHCII binding (**C**) AF3 models of epitope B4 Nb lead B115 which interact with alpha helix 2 and the N-terminal beta barrel domain to block TCR binding. (**D-F**) Epitope B1 Nb AF3 models, showing CDR interaction with alpha helix 6 and beta strand 11 for potential MHCII competition.

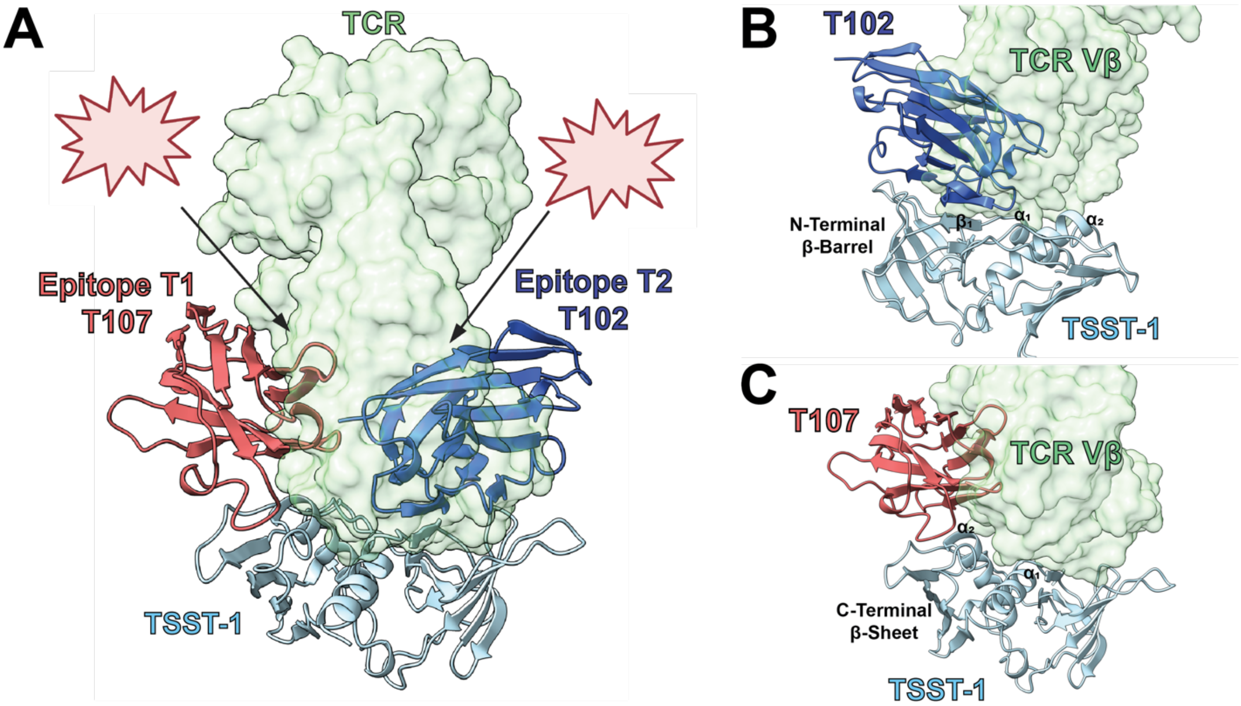


**Supplementary Figure 6. AlphaFold 3 (AF3) models of anti-TSST-1 Nbs (A**) All high confidence AF3 models of TSST-1 Nbs (iPTM>0.8) overlayed on TSST-1. Structures of TSST-1 co-complexed with TCR V beta (PDB: 2IJ0) and the fully assembled TCR (PDB: 4C56) were superimposed to view TCR competition. (**B**) AF3 models of epitope T2 Nb T102 which interact with beta strand 1, alpha helix 1, and alpha helix 2 and the N-terminal beta sheet of TSST-1 to block TCR binding (**C**) AF3 models of epitope T1 Nb T107, which interacts with helix 2 and the C-terminal beta sheet of TSST-1 to block TCR binding.

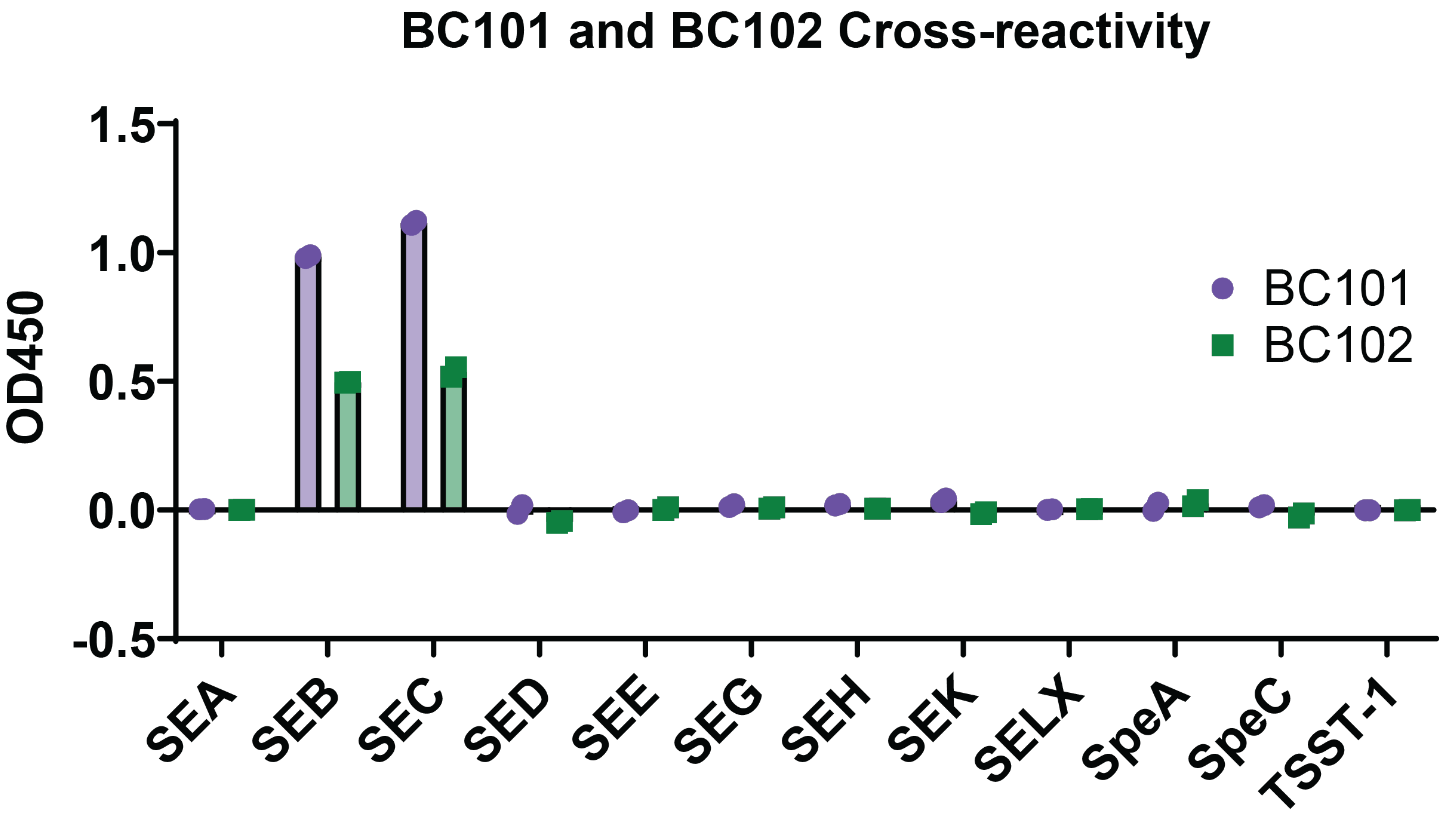


**Supplementary Figure 7.** Single saturating concentration ELISA binding screen of cross-reactive Nbs against select SAgs. SAgs coated at 10µg/ml and all Nbs assessed at 1µM Nb using HRP conjugated anti-T7 polyclonal antibody. All experiments conducted in duplicate.

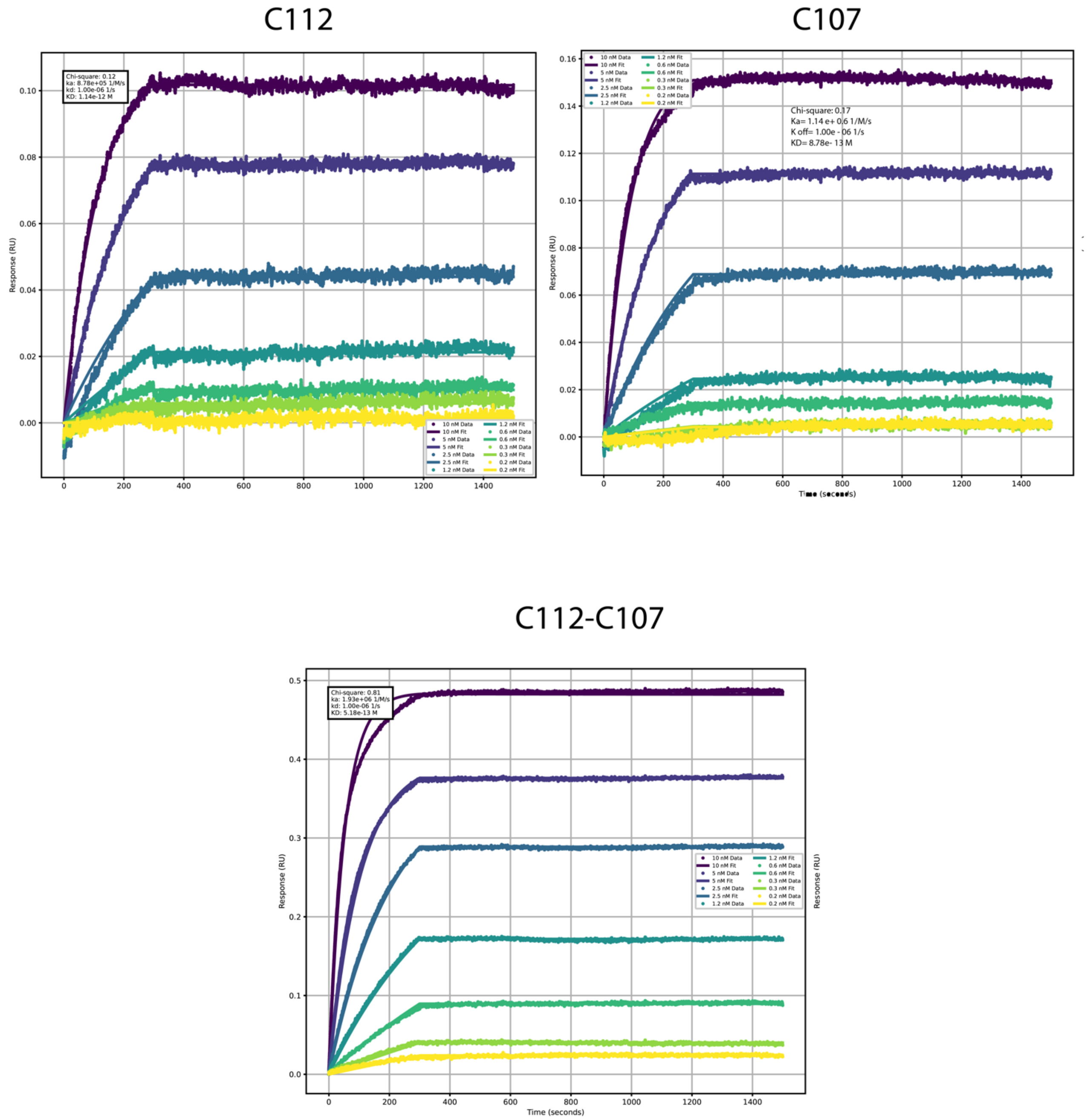


**Supplemental Figure 8**. BioLayer Interferometry (BLI) analysis of three Nb constructs (C112, C107 and the biparatopic C112-C107) for SEC binding.

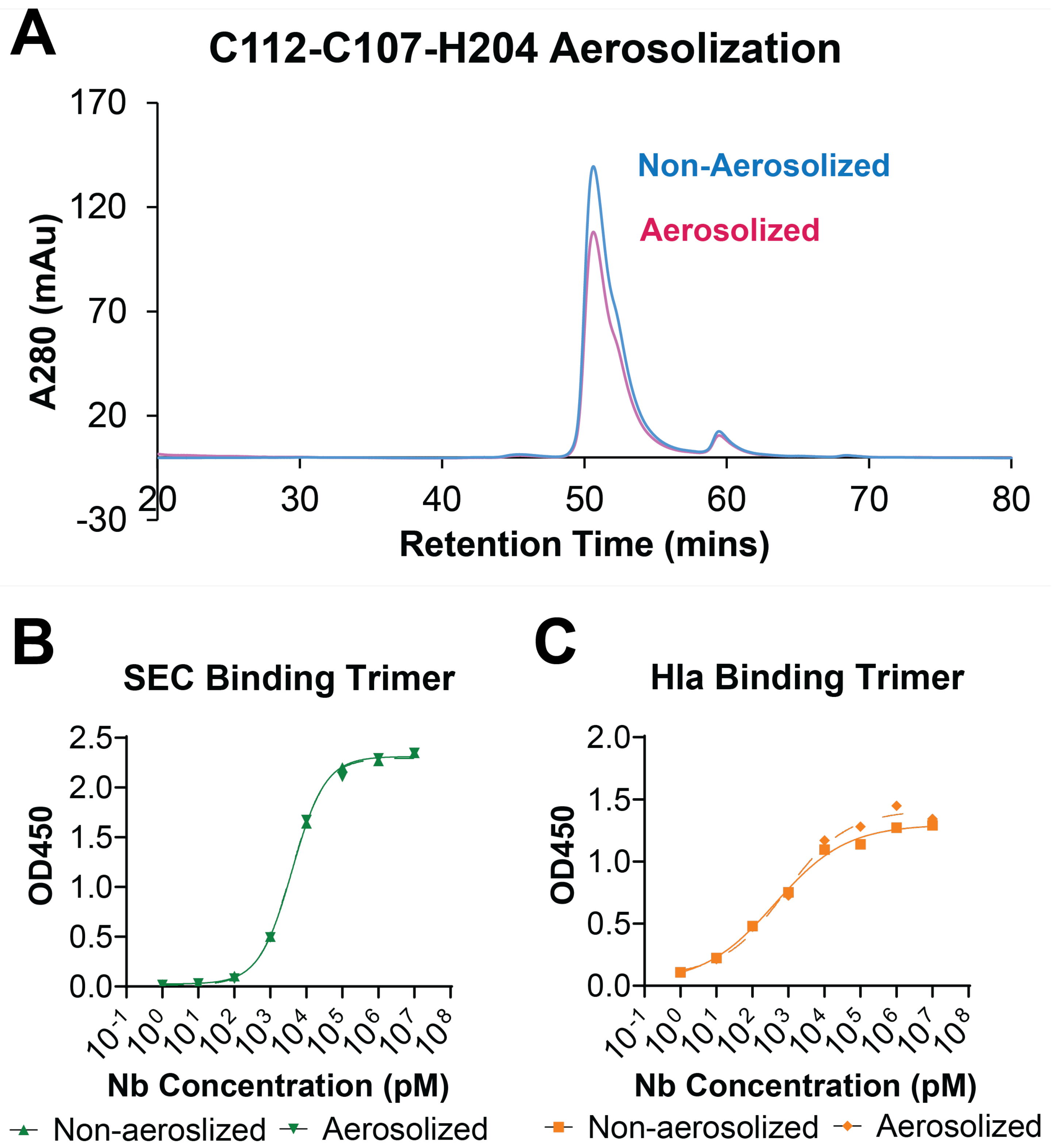


**Supplementary Figure 9.** (**A**) Size exclusion chromatography of C112-C107-H204 before and after aerosolization. (**B**-**C**) ELISA binding activities of C112-C107-H204 against SEB (**B**) and Hla (**C**) before and after aerosolization (n=1).

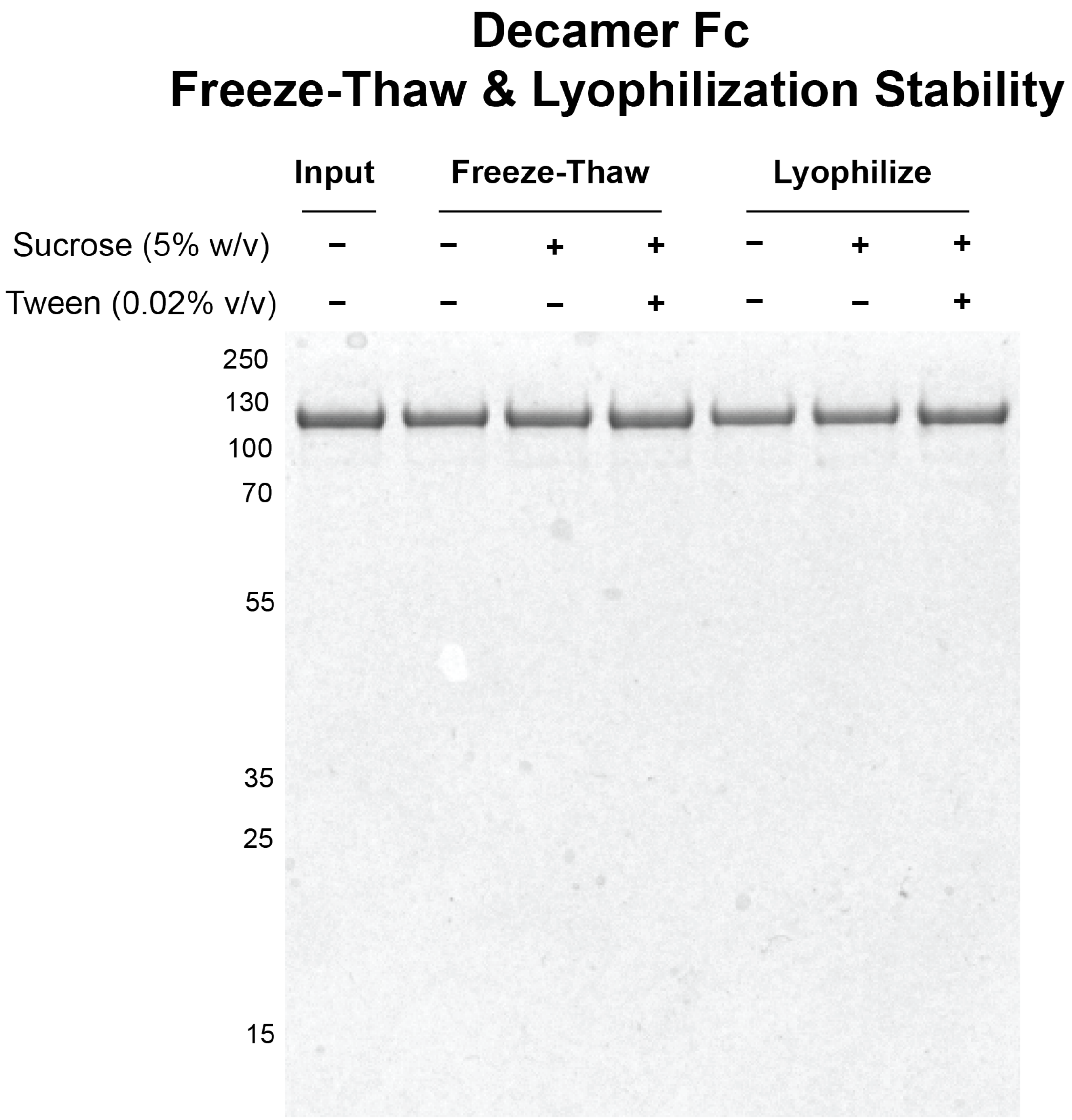


**Supplementary Figure 10.** SDS-PAGE analysis of Decameric Nb Fc YTE construct with sucrose cryoprotectant and Tween detergent after freeze-thaw (lanes 2-4) and lyophilization-reconstitution (lanes 5-7) compared to input (lane 1).

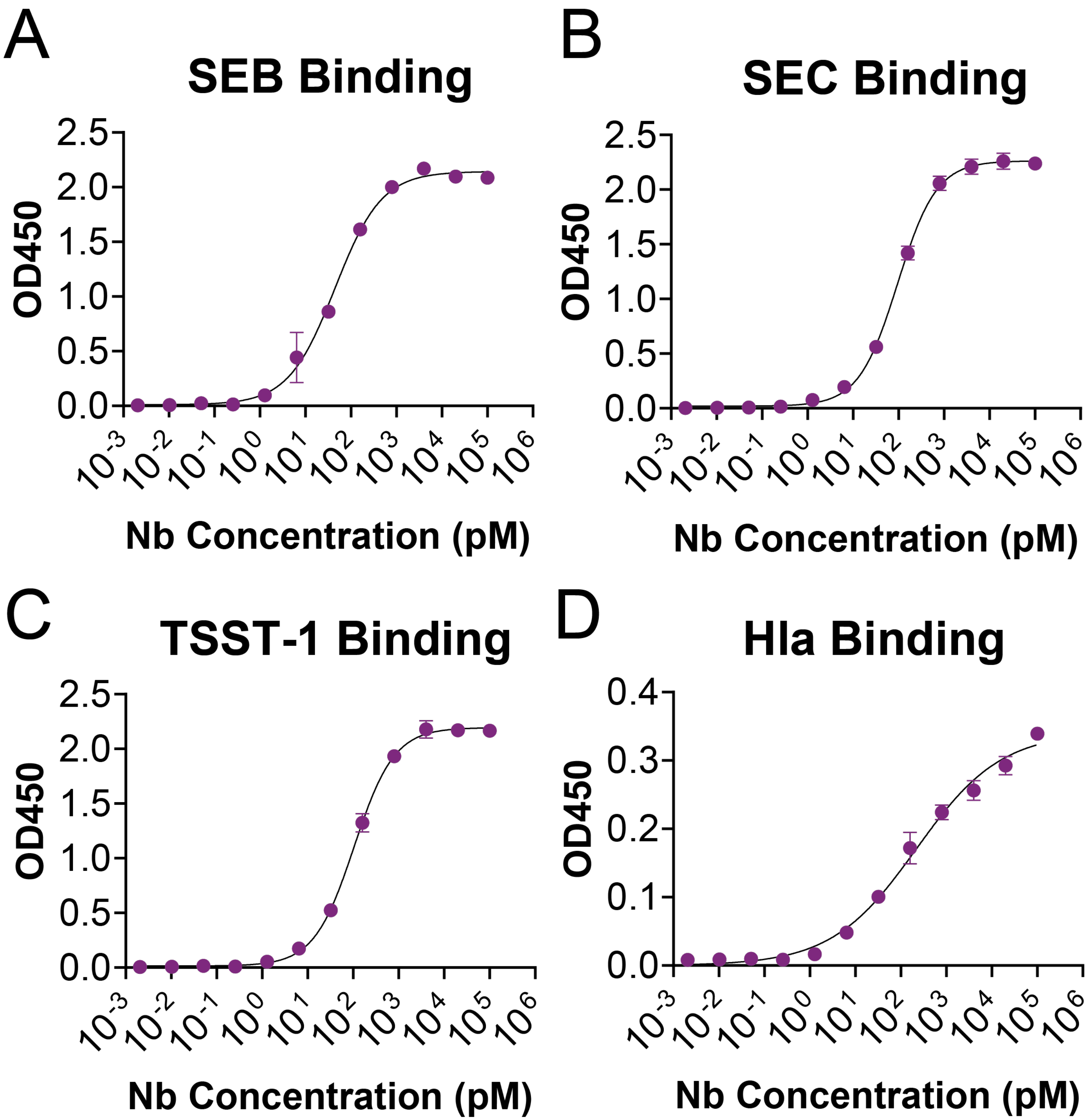


**Supplementary Figure 11.** ELISA binding activity of decameric Nb Fc YTE fusion construct against (**A**) SEB, (B) SEC, (C) TSST-1, and (D) Hla. All binding assessments done in triplicate and fitted according to 4PL regression analyses.

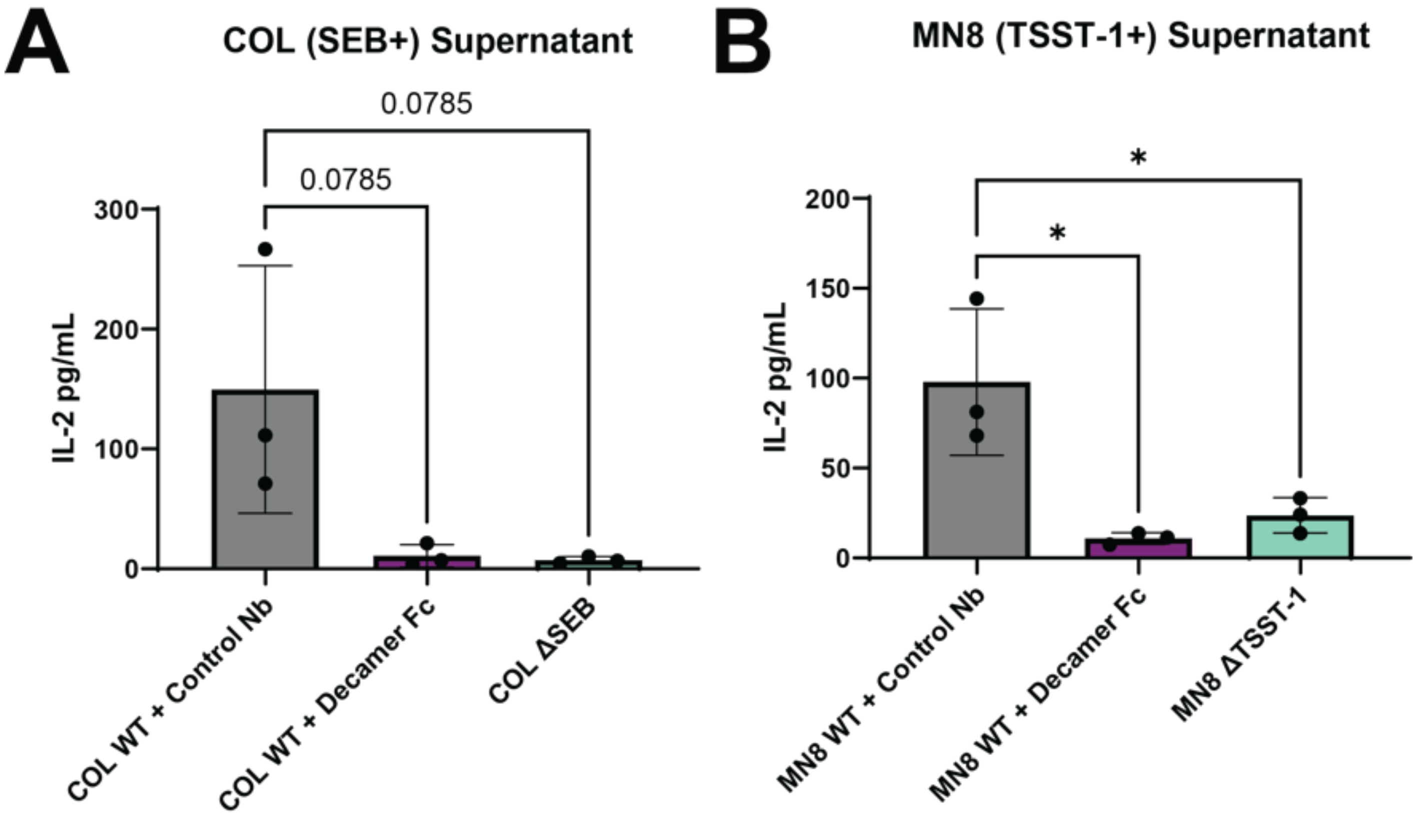


**Supplementary Figure 12.** PBMC stimulation assays to assess Decameric Nb Fc neutralization of overnight supernatants of (A) COL strain and (B) MN8 strain. Experiments conducted in triplicate and analyzed using one way ANOVA analyses (* is $p<0.05$).
